# DNA barcode developement based on chloroplast and ITS genes for species identification of endangered and threated species of Western Ghats, India

**DOI:** 10.1101/2024.06.04.597498

**Authors:** Tanzeem Fatima, MN Srividya, Raj Kishore Singh

## Abstract

Accurate identification is crucial for conserving species, especially in regions such as the Western Ghats, where trade poses a significant threat to endangered and threatened forest species. Traditional morphology-based identification can be challenging and time-consuming, leading to inaccuracies, especially with similar-looking species or dried specimens. Therefore, DNA barcoding offers a potent solution for precise species identification to address illicit trade and address impactful conservation measures. DNA barcoding is a taxonomic technique that uses standardized short DNA sequences to differentiate and classify species. This approach is especially valuable when morphological characteristics alone are insufficient for accurate species identification. In this study, we focused on the development of a DNA barcoding system for the efficient and accurate identification of threatened and endangered important forest species of Western Ghats Karnataka. To develop the DNA barcoding system, a multilocus approach utilizing sixteen standard DNA barcoding markers was used. A total of 47 threatened and endangered forest species from the Western Ghats were selected for this study. Using a larger number of markers to develop DNA barcodes led to the most precise species identification rates. Moreover, the wide availability of DNA barcode databases allows for quick and accurate species identification. In our study, we observed the highest amplification rates for rbcL1 (40 species), psbtrnH2 (36 species), and PsbA-trnH1 (33 species). DNA amplification varied from 11.76% to 94.11%. Notably, the highest DNA amplification rates were detected for *A. wightii* (94.11%) and *A. hondala (*92.34%), both of which belong to the Arecaceae and Passifloraceae families, respectively. Sequencing success rates ranged from 37.5% to 100%. This study will aid in the development of a database of available threatened forest species in western Ghats Karnataka and other regions.

## Introduction

The identification of forestry species plays a crucial role in biodiversity conservation, particularly in regions such as the Western Ghats Karnataka, where plant biodiversity hotspots face threats from habitat fragmentation, overexploitation, and human activities. The preservation of threatened and endangered plant species is essential for biodiversity conservation efforts (Kress et al. 2005). However, traditional methods of plant identification are facing challenges due to declining taxonomic expertise and the lack of precise tools to distinguish between various plant forms, such as seeds, plant parts, seedlings, and herbal adulterants (Saddhe and Kumar 2018). Recognizing commercial plant materials can be particularly difficult because they are often traded in dry or processed forms and lack the necessary morphological features for accurate identification by retailers and customers (Ghorbani et al 2017). Taxonomic identification using macro- and micromorphological techniques is laborious, prone to errors, and requires expertise and reliable resources. DNA barcoding has emerged as a promising method for accurately identifying plant species, offering distinct advantages over traditional morphological identification techniques (Saddhe and Kumar 2018). This molecular marker-based approach allows for rapid screening of DNA variation, resulting in increased taxonomic resolution (Fatima et al 2019). It has proven particularly effective for species with ambiguous morphological features (Ghorbani 2017). Despite extensive research on DNA barcoding for trees and plant species, a significant number of important species, particularly those found in the Western Ghats, still lack DNA barcode sequences for molecular identification. In the literature, it has been noted that barcoding and DNA sequences are accessible for certain species. However, a significant portion of species currently lack available DNA barcodes, highlighting a gap in our knowledge and resources within this domain. Our study aimed to address this gap by providing DNA barcodes for threatened forest species, thereby supporting conservation efforts in western Ghats Karnataka. To ensure accuracy, researchers often employ a multilocus approach utilizing various DNA markers (Saddhe and Kumar 2018; Abubakar et al 2017; Li et al 2011). We evaluated the efficacy of sixteen sets of primers targeting different regions in the nuclear and plastid genomes of plants, including commonly used markers such as matK, rbcL, trnH-psbA, Ar matk (Cuenoud et al. 2002), rpoB, rpoC (Kurian et al 2020, Sass et al 2007), rpoB-trnCGAR (Anerao et al 2021), psbZ-trnfM (Kurian et al 2020; Scarcelli et al 2011), trnHpsbA (Kurian et al 2020; Kress et al 2005), psbK (Kurian et al 2020; Lee et al 2007), rbcL (Fatima et al. 2019), PRK (Lewis and Doyle 2002), RPB2, ITS2, ATP (Kurian et al 2020), ndhF-rpl32 (Sass et al. 2007), MatK2 (Cuenoud P et al 2002), psbA-trnH (Chandrasekara et al 2021, Sang et al 1997), Ar rbcL (Jeanson M L et al.2011), and DI ITS (Fatima et al 2019)

In our study, our goal was to create a robust DNA barcoding system capable of accurately identifying 47 endangered and threatened forest species from 21 different families*, namely,* Fabaceae, Lauraceae, Santalaceae, Balsaminaceae, Myristicaceae, Ebenaceae, Annonaceae, Rubiaceae, Arecaceae, Dipterocarpaceae, Cluciaceae, Calophyllaceae, Myrtaceae, Malvaceae, Apiaceae, Apocynaceae, Asteraceae, Passifloraceae, Sapotaceae, Chrysobalanaceae, Cycadaceae and Anacardiaceae, providing important insights into their conservation and management (Table 1) of forest species in western Ghats Karnataka. The results of this study contribute to the development of a molecular database containing information on threatened forest species not only in western Ghats but also across other regions. Such a database can significantly aid conservation efforts by enabling accurate identification and monitoring of endangered species and threatened forest species.

**Table 1.**
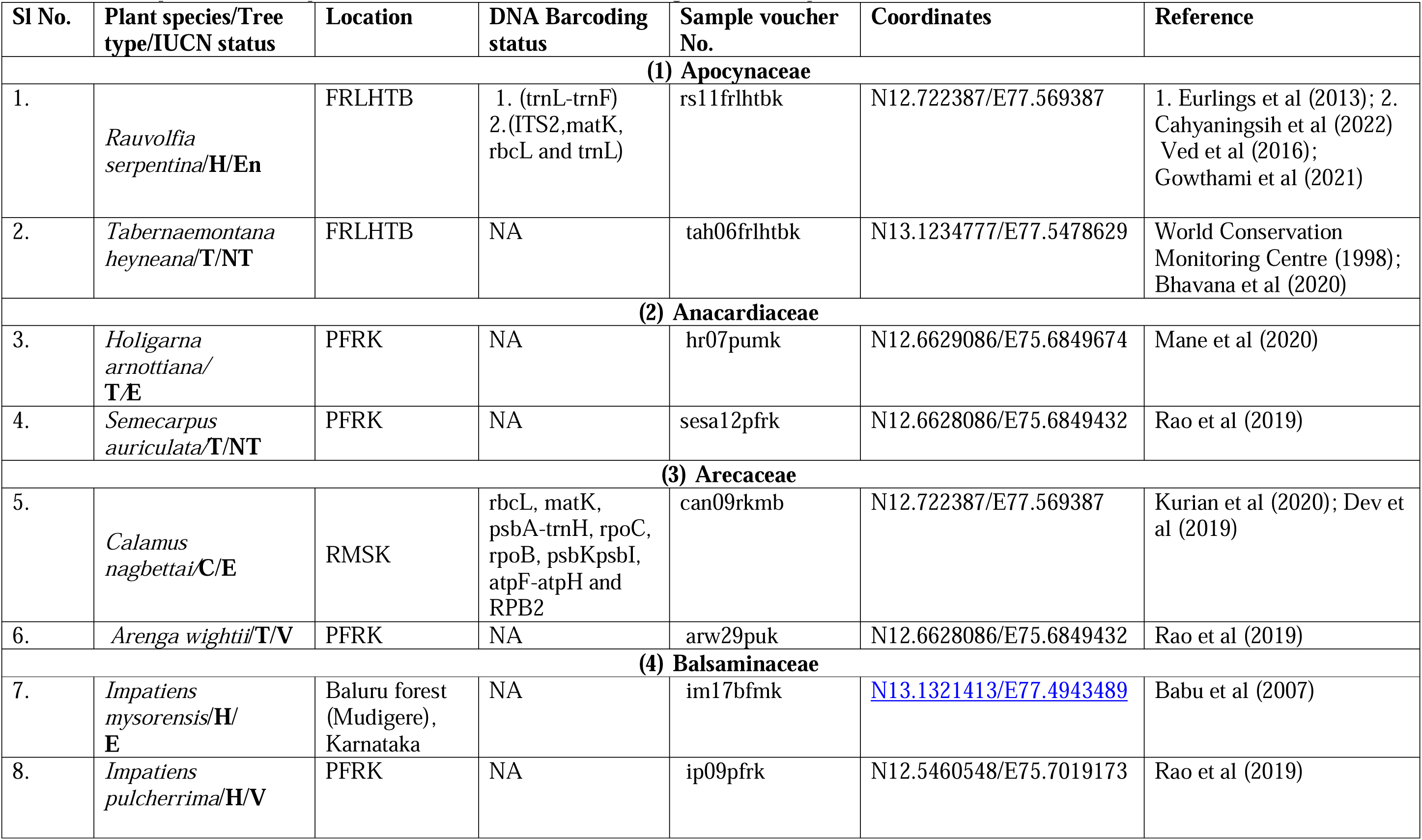
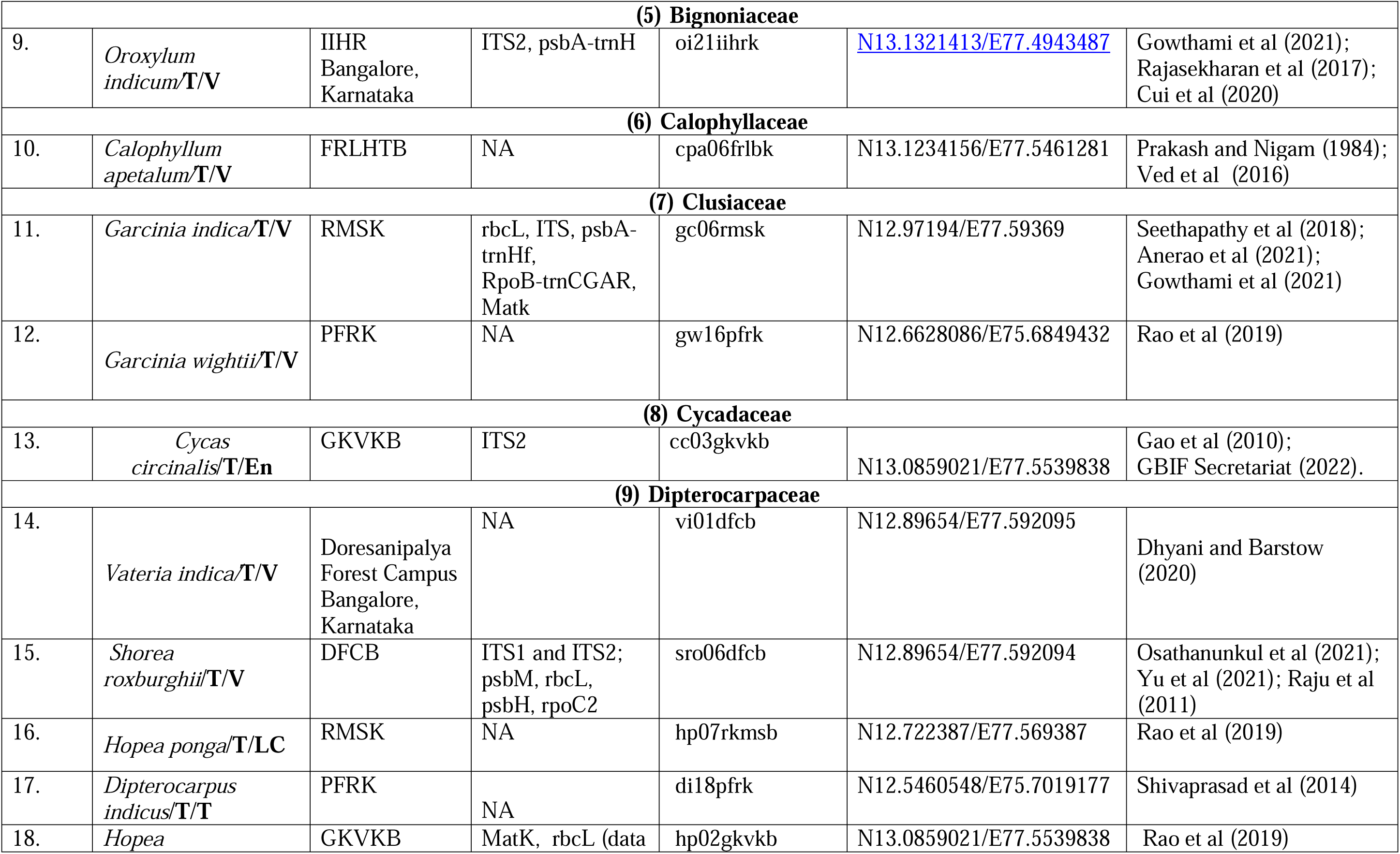
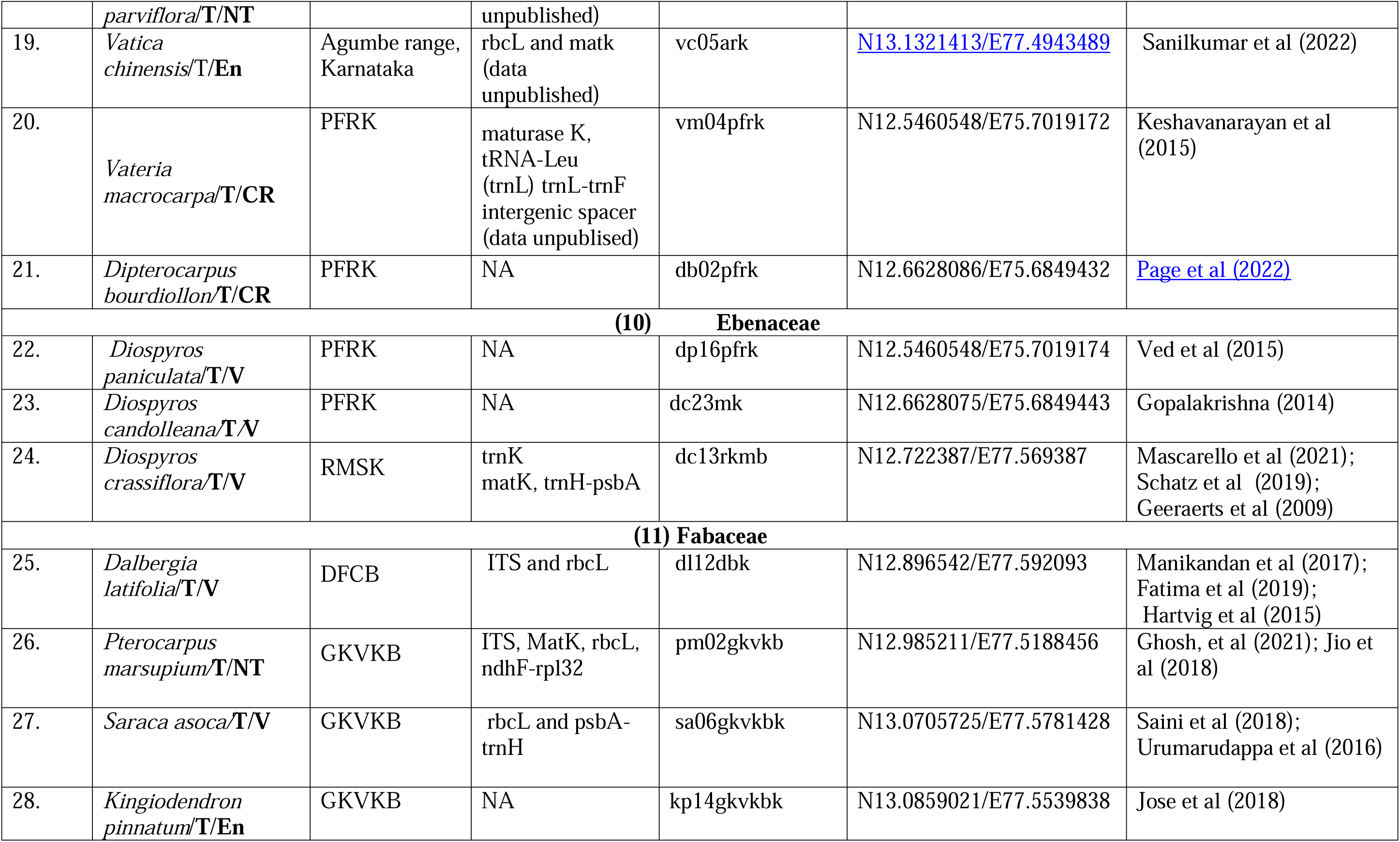
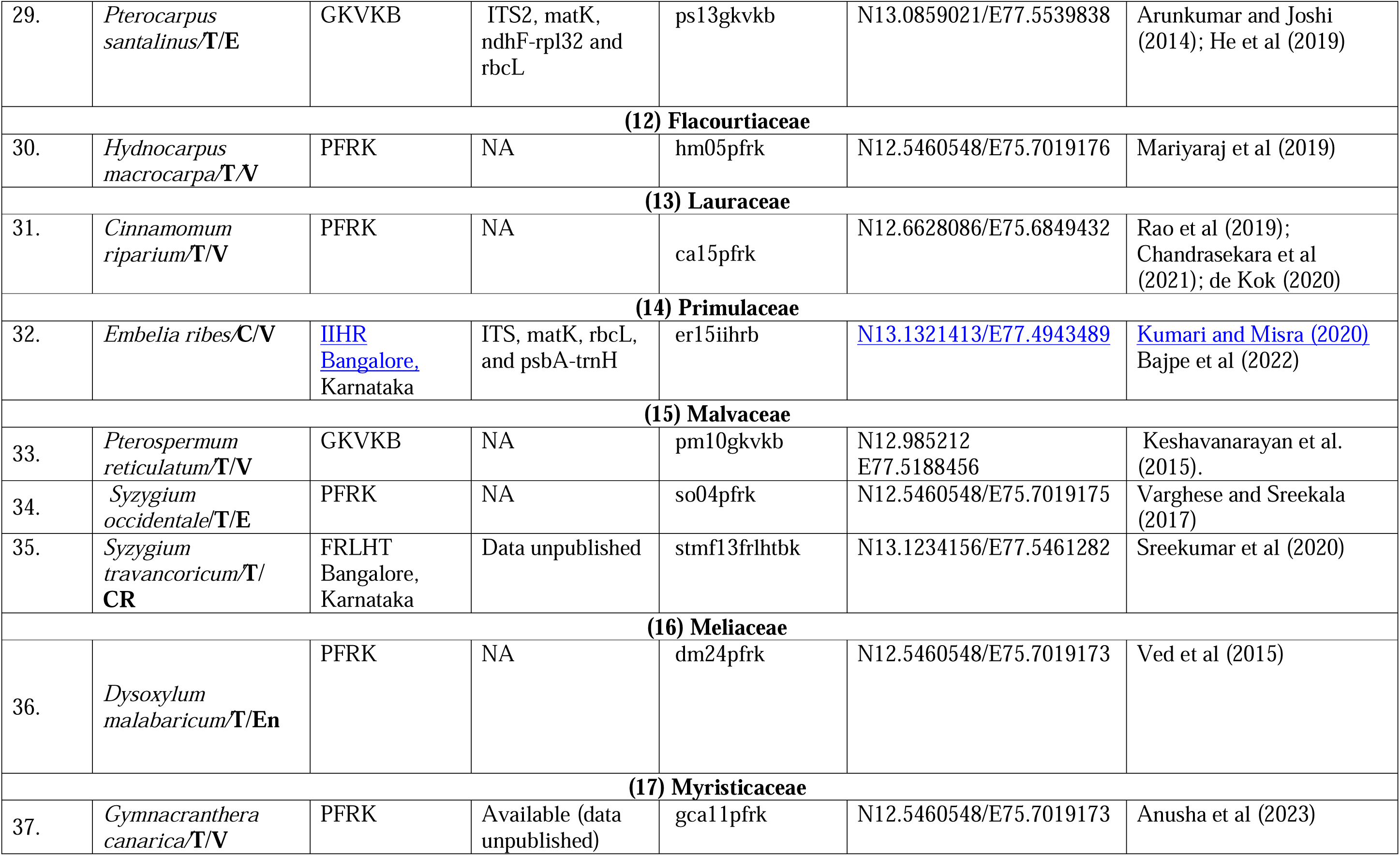
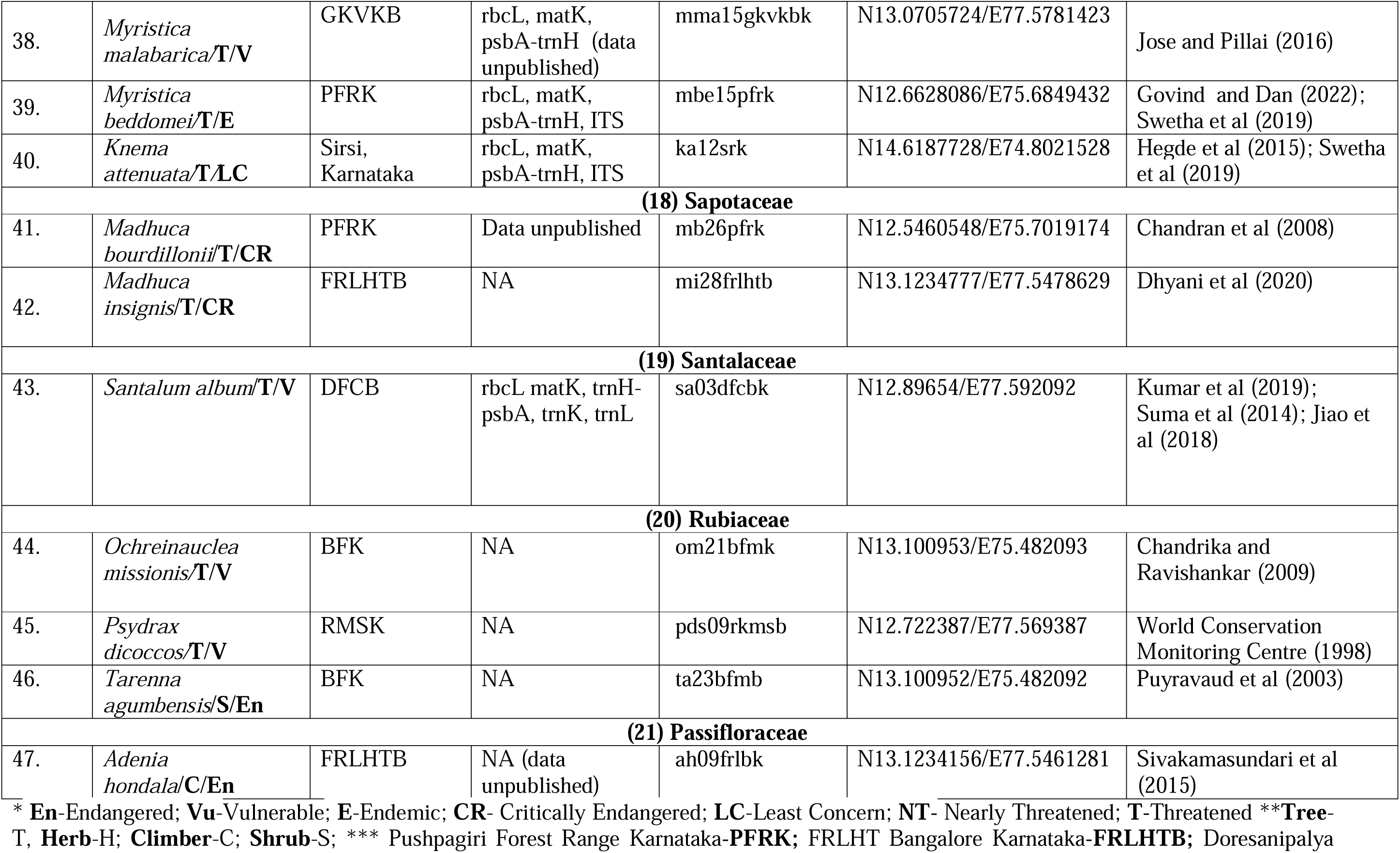
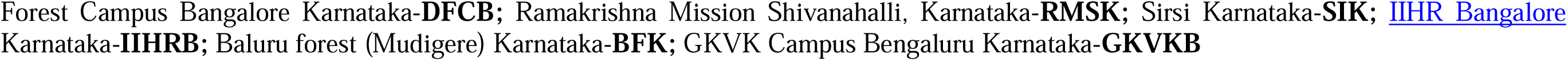
Description of the sample collection of Threatened and Endangered forestr Species of Karnataka.

## Methodology

### Study area

The sampling site for this study consisted of various locations across Karnataka. The samples of selected species were collected from the regions of Bengaluru, Sirsi, Madikeri, Agumbe, and Chikkamagaluru. These locations were selected based on their availability and relevance to the study of threatened and endangered plant species in the region. Samples from a total of 47 different threatened and endangered species were collected, each of which was carefully chosen to represent the overall biodiversity of the area. To maintain the accuracy of the study, the GPS coordinates of each sampling site were recorded during the fieldwork. This information was used to map the distribution of the selected species across the various locations in Karnataka (Fig. 1).

**Fig. 1.**
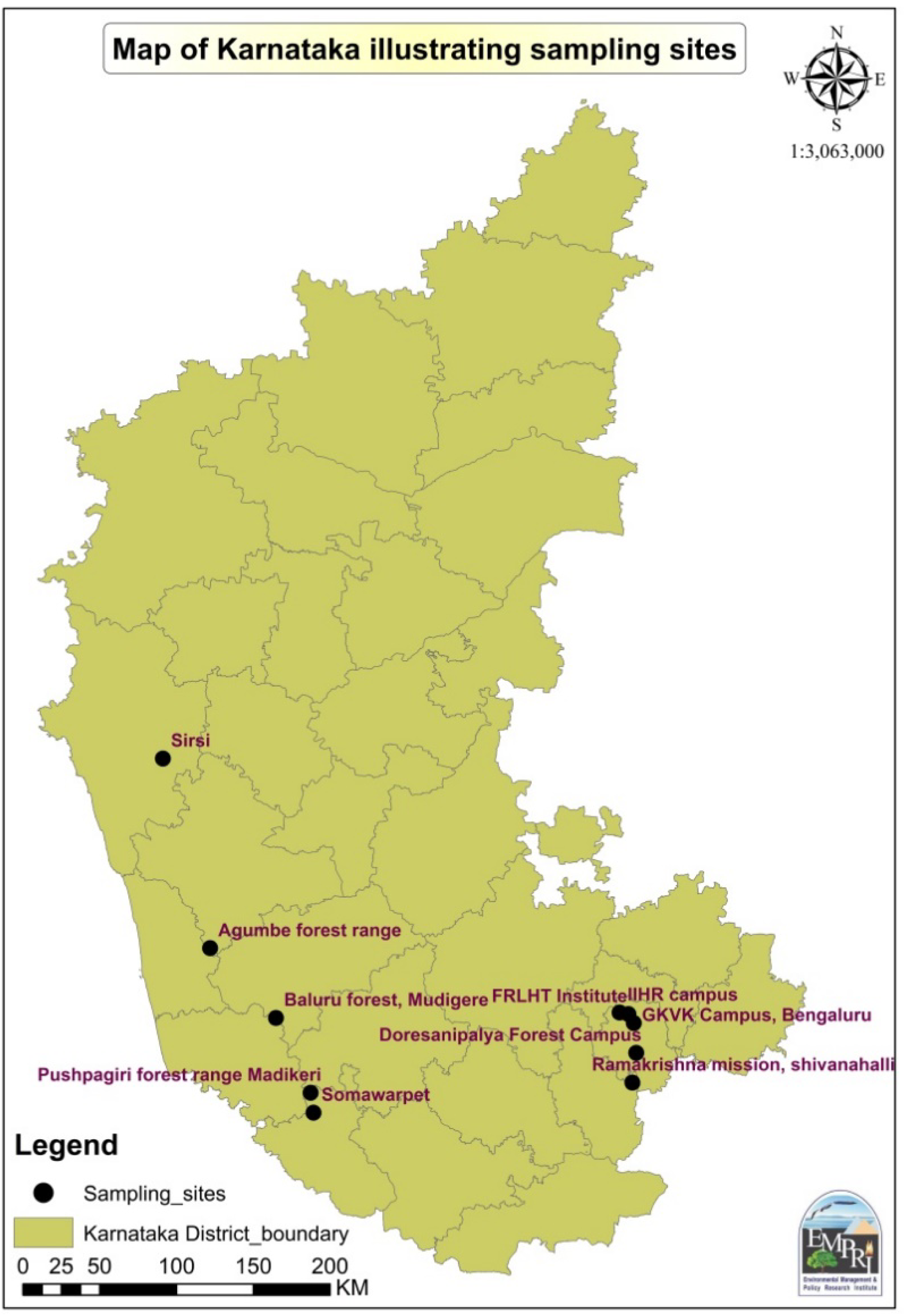
Map of Karnataka showing the sampling sites.

Details about the selected plant species, sample locations, GPS coordinates, and altitudes were recorded (Table 1), providing a comprehensive overview of the environmental conditions present at each location. This information can be used to better understand the factors that contribute to the survival of these threatened and endangered species in the region.

The data collected from these locations can provide valuable insights into the conservation of these species and their habitats.

### 2. Sample collection

The sample collection for this study involved the collection of leaf samples from 47 selected threatened and endangered species from various parts and forests of the western Ghats. To ensure a representative sample, five to ten samples (five to ten leaves) of each species were selected.

Leaf sampling was carried out by carefully selecting healthy leaves from each plant, taking care not to damage the plant. The leaf samples were then placed in a plastic cover to prevent any damage or contamination during transport.

To prepare the samples for further analysis, the collected samples were cleaned and left to dry for a period of at least seven to fourteen days, allowing the moisture content of the samples to be eliminated. This process helps to preserve the samples for further analysis and helps to prevent the growth of fungi and other microorganisms that can degrade the samples.

In addition to sample preservation, herbarium sheets were prepared for each species. The process of preparing the herbarium sheets involved pressing and drying the collected plant specimens onto acid-free herbarium sheets. The specimens were then labeled with information such as the species name, collection date, location, and collector’s name. This ensures that the samples can be identified and referenced in the future, even after the plant material has deteriorated.

### 3. Genomic DNA isolation and quantification

Total genomic DNA was isolated from the dried leaves by using a modified CTAB protocol (Fatima et al. 2018, Fatima et al. 2019). The extracted DNA was quantified, and to confirm the DNA’s suitability for sequencing and barcoding, the samples were run on a 1% agarose gel stained with 4 μL of 0.3% ethidium bromide (Fatima et al. 2019).

### 4. PCR amplification and purification of amplified PCR products

Sixteen sets of primers were selected from the literature for the current DNA barcoding study. The details of the primer sequences are shown in Table 2. A reaction volume of 13 μL was used for DNA amplification, containing 1.5 μL of genomic DNA (30 ng/μL), 2 μL of each forward and reverse primer (10 mM), 1.5 μL of PCR buffer, 1.5 μL of dNTPs (10 mM), 1.5 μL of MgCl_2_, 0.3 μL of Taq polymerase (3 U/μL) (Bangalore genus), and 4.2 μL of double-distilled water. The amplification cycle consisted of an initial denaturation step for 3 min at 94 °C, followed by 40 cycles of 1 min at 60–61 °C and 1 min at 72 °C and a final extension step for 10 min at 72 °C (Williams et al. 1990).

**Table 2.**
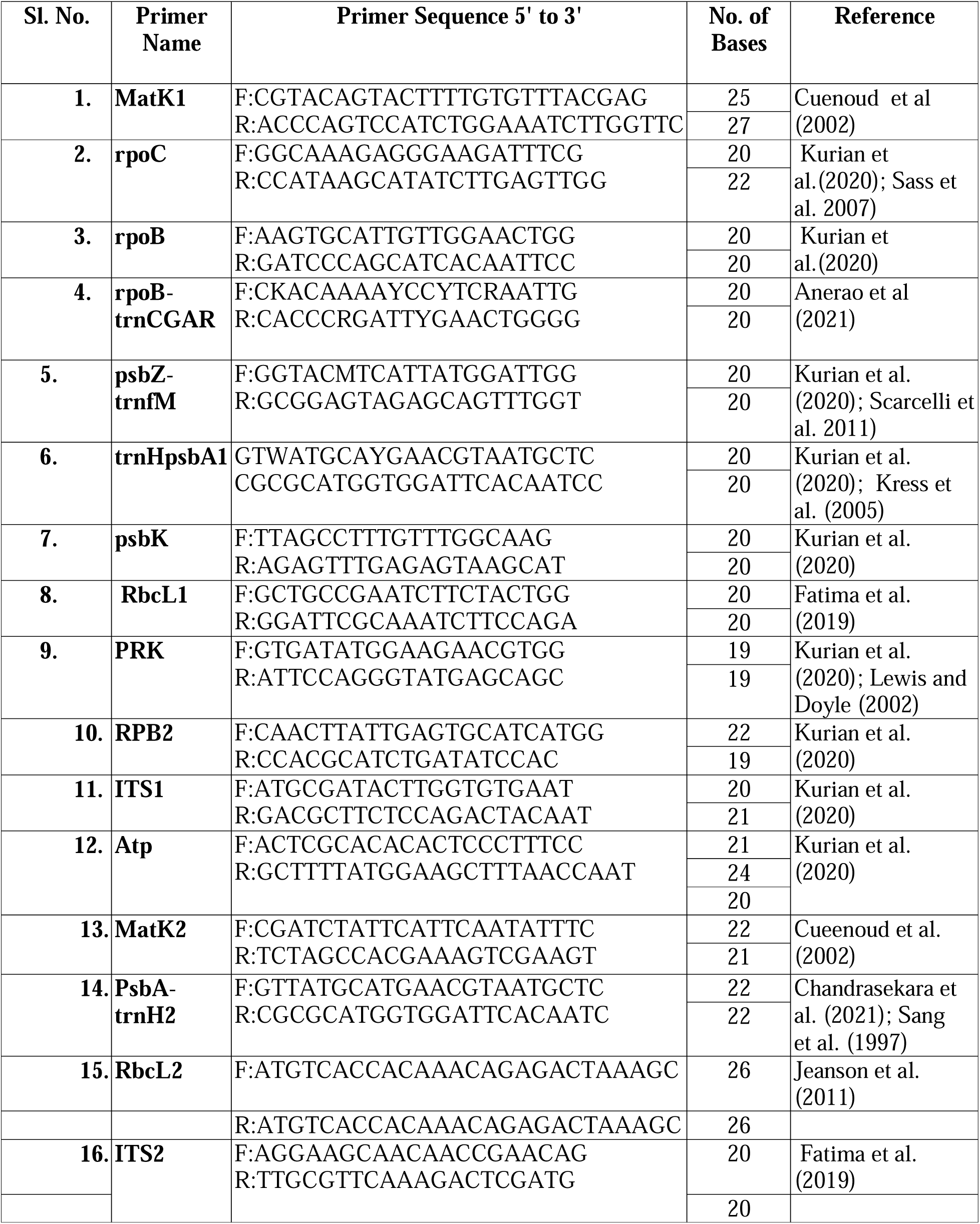
Primers used for DNA barcoding.

### 5. DNA sequencing and DNA barcode generation

The purified PCR products were subjected to Sanger sequencing at Eurofins Genomics India Pvt. The primers and barcodes for each sequence were developed by using Bio-Rad online software. The developed sequences were submitted to the NCBI Sequence Data Submission Bank (http://www.ncbi.nlm.nih.gov/gen-bank/) to obtain accession numbers. The nucleotide database of the US National Centre for Biotechnology Information (NCBI, www.ncbi.nlm.nih.gov) was searched for homologs of related genes.

### Testing DNA barcode accuracy

Before evaluating the identification success of the selected DNA barcode regions, we used the BLAST tool (https://blast.ncbi.nlm.nih.gov/Blast.cgi) to compare our sequences with those submitted to GenBank, which will be accessed on February 10, 2025 (https://www.ncbi.nlm.nih.gov/genbank/), with the aim of confirming the molecular level of identification of selected species.

## Results and Discussion

The 47 endangered, vulnerable, and threatened forest species selected for the project represent diverse plant families. Among these families, Dipterocarpaceae had the greatest number of species, totaling eight. Fabaceae had five species, and Myristicaceae had four species (Fig. 2; Table 1).

**Fig 2.**
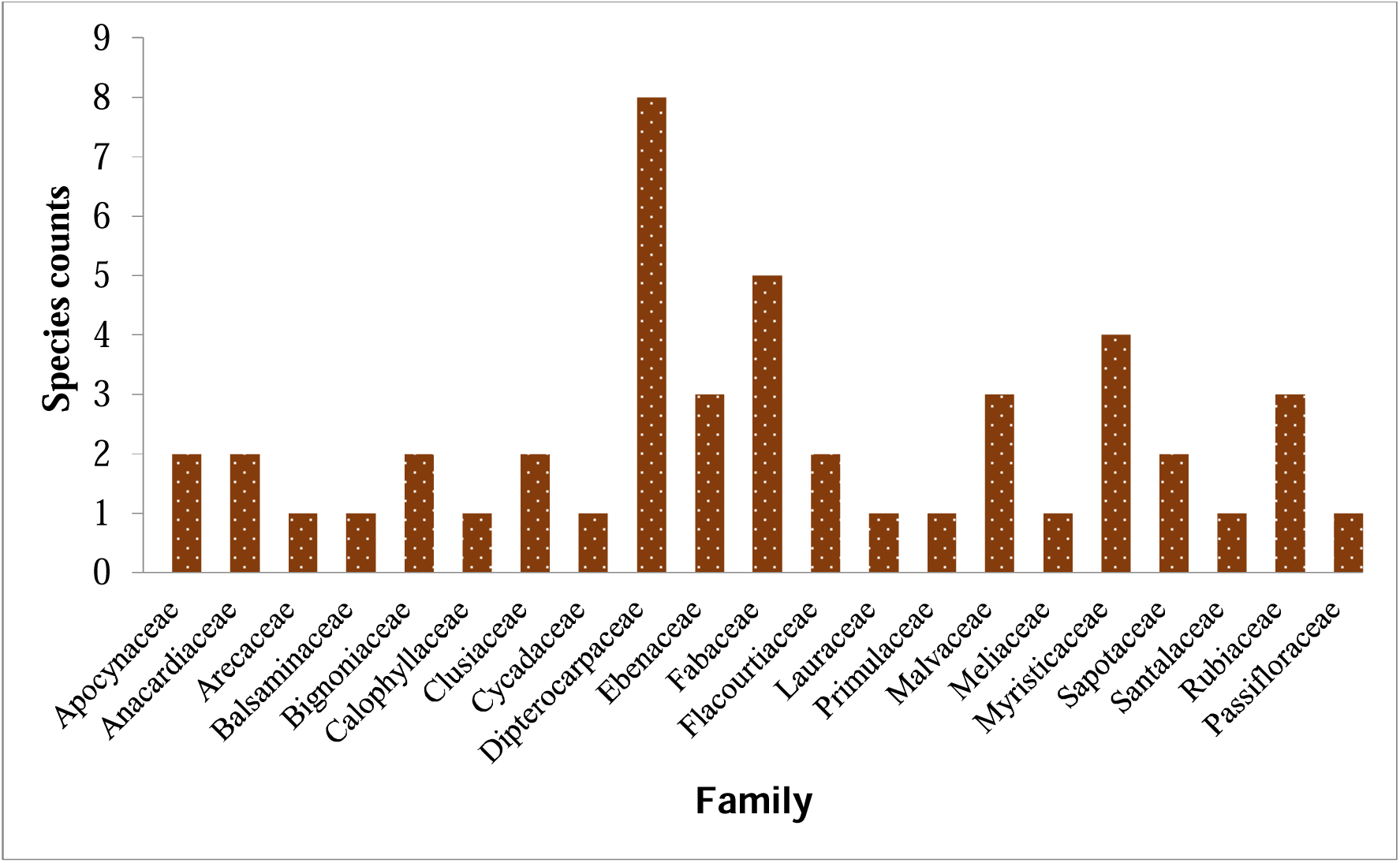
Species count based on the family.

### Frequency of selected marker amplification

In this study, a total of sixteen DNA barcoding markers were selected from various literature references (Table 2). When examining these markers individually, some were notably more effective at identifying species than others were. For instance, the RbcL1 marker emerged prominently with a high success rate of 37% in species identification, closely followed by PsbA-trnH2 and PsbK, which achieved success rates ranging from 36% to 32%. However, some Barcoding markers, such as MatK2 and ITS2, exhibited lower amplification rates (Fig 3).

**Fig 3.**
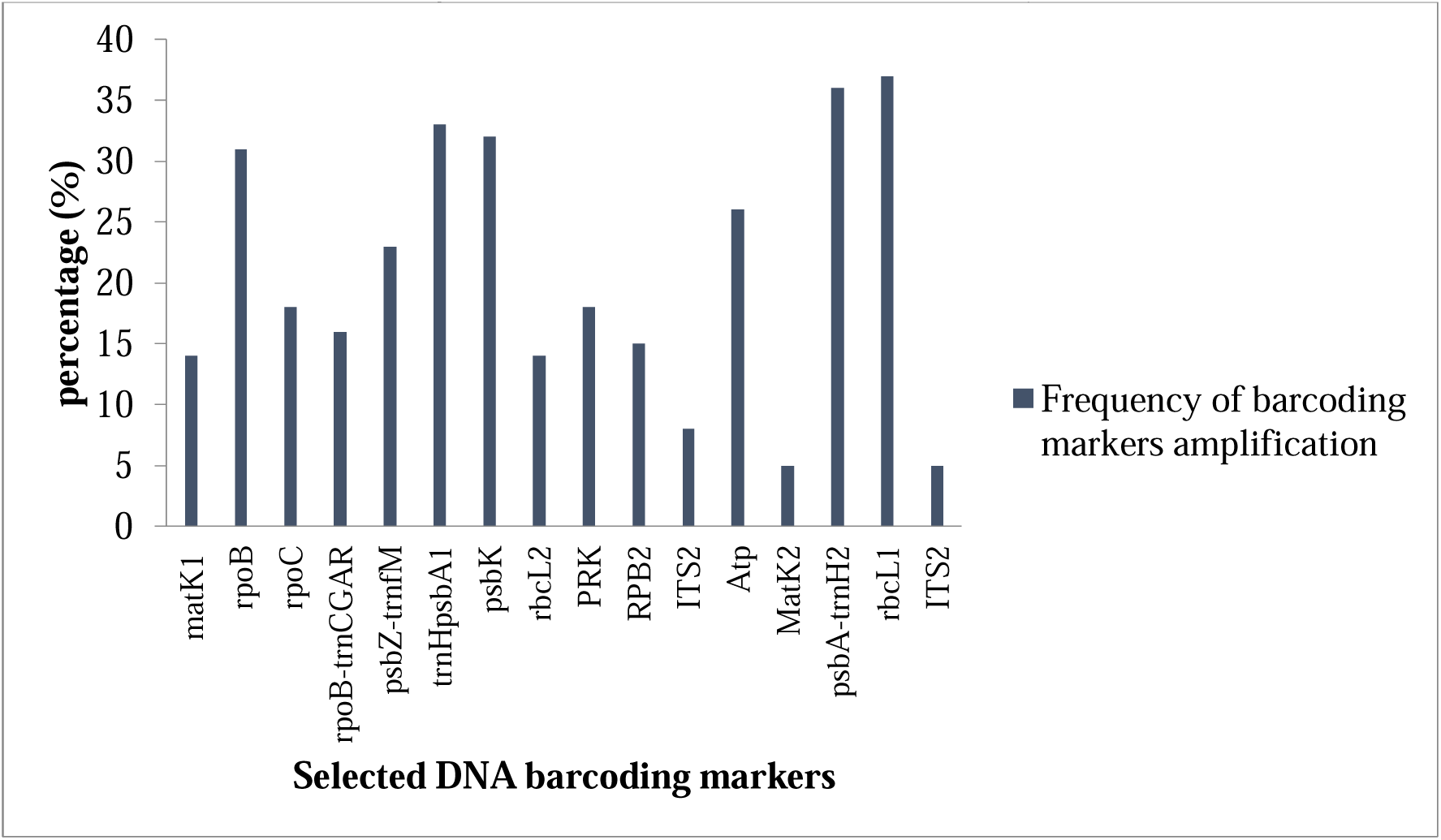
Amplification frequency of the selected DNA barcoding markers.

### DNA barcoding marker amplification and sequencing across selected families and their respective species

The present investigation listed 21 families according to the chosen species (Table 1). All of these samples were amplified using selected DNA barcoding markers. While amplification was detected in all species, it was limited to only a few markers in certain instances. DNA amplification ranged from 11.76% to 94.11%. The highest DNA amplification rates were observed for *A. wightii* (94.11%) and *A. hondala* (92.34%), which belong to the Arecaceae and Passifloraceae families, respectively. Conversely, the lowest amplification success rate (30.76%) was noted for four species, namely, *T. heyneana, V. indica, P. marsupium* and *M. insignis*. The sequencing success rates ranged from 37.5% to 100%, with twenty out of the 47 selected species achieving a 100% sequencing success rate (Fig 4). *The* Apocynaceae family includes species such as *R. serpentina* and *T. heyneana*. Both species were analyzed using DNA amplification techniques, with common markers such as RpoB, RbcL1 and Atp. Additional markers, such as PsbZ-trnfM, trnHpsbA1, and PsbA-trnH2, were detected in *R. serpentina*, while *T. heyneana* was amplified solely with the PsbK marker. The collective amplification rate was 46.15± 30.76%, with a subsequent sequencing rate of 100% observed across the respective species within the family. The Anacardiaceae family comprises species such as *H. arnottiana* and *S. auriculata*. DNA amplification techniques were applied to both species utilizing common markers such as PsbK, PsbA-trnH2, and RbcL1. *H. arnottiana* was amplified with the trnHpsbA1, RPB2, and Atp markers, while *S. auriculata* was amplified with the MatK1, RpoC, and RpoB-trnCGAR markers. The collective amplification and sequencing rates were as follows: *H. arnottiana* (76.92 ± 100%) and *S. auriculata* (52.94 ± 88.9%). In the Arecaceae family, the analysis of two species, *C. nagbettai* and *A. wightii*, revealed intriguing findings. Both species were amplified using common markers such as RpoB, PsbZ-trnfM, and RbcL1. Notably, *A. wightii* exhibited additional amplification with markers such as MatK2, RpoC, and RbcL2, revealing a broader genetic profile. The amplification rates for each species were distinct: *C. nagbettai* displayed amplification and sequencing rates of 87.65% ± 100%, while *A. wightii* showed a robust amplification rate of 94.11% ± 87.5%. In the context of the Balsaminaceae, the investigation included *I. mysorensis* and *I. pulcherrima*. Both species were amplified using common markers such as trnHpsb1, PsbA-trnH2, and PsbK. Notably, *I. mysorensis* displayed additional amplification with RpoB, while *I. pulcherrima* exhibited amplification with markers such as RbcL2 and MatK1 (Table 3). The amplification and sequencing rates for each species were as follows: *I. mysorensis* (46.15 ± 100) and *I. pulcherrima* (64.7 ± 90.9). Within the Bignoniaceae family, investigations have focused on *O. indicum*. This species exhibited amplification with multiple barcoding markers, including MatK1, RpoC, and RpoB (Table 3). The amplification and sequencing rates observed for *O. indicum* were 64.7 ± 100%, respectively. Within the Calophyllaceae family, *C. apetalum* is a solitary member. Notably, *C. apetalum* demonstrated amplification and sequencing rates of 61.5 ± 87.5%. This species shows amplification and sequencing patterns involving markers such as RpoB, PsbZ-trnfM, and trnHpsbA1. In the Clusiaceae family, DNA barcoding analysis was conducted on two species, namely, *G. indica* and *G. wightii*. While *G. indica* exhibited amplification with only two specific barcode markers, namely, PsbZ-trnfM and ITS2, Garcinia wightii displayed a more extensive amplification profile. Twelve barcoding markers, namely, MatK1, RpoC, RpoB, RpoB-trnCGAR, PsbZ-trnfM, trnHpsbA1, PsbK, RbcL2, RPB2, PsbA-trnH2, RbcL1, and Atp, were successfully captured from G. wightii. The amplification and sequencing rates observed for *G. indica* and *G. wightii* were 38.46 ± 40.0% and 76.47 ± 92.3%, respectively. In the Cycadaceae family, DNA barcoding analysis was performed on *C. circinalis*, which revealed the amplification of four distinct DNA barcode markers. These markers, namely, RpoB-trnCGAR, PsbZ-trnfM, PRK, and Atp, contributed to the generation of unique DNA barcodes. The observed amplification and sequencing rates for *C. circinalis* were 53.84 ± 57.1%. A comprehensive investigation revealed various species of the Dipterocarpaceae family, including *V. indica, S. roxburghii, H. ponga, D. indicus, H. parviflora, V. chinensis, V. macrocarpa* and *D. bourdiollon*. Each species exhibited specific amplification patterns during genetic analysis. *V. indica* was amplified with four barcoding markers: trnHpsbA1, PsbK, PsbA-trnH2, and RbcL1. The amplification of the RpoB, trnHpsbA1, PsbK, RPB2, PsbA-trnH2, RbcL1, and ITS2 markers in S. roxburghii is shown. *H. ponga* displayed amplification of RpoB-trnCGAR, trnHpsbA1, PsbK, RPB2, PsbA-trnH2, RbcL1, and ITS2. *D. indicus* was amplified with the markers MatK1, RpoC, RpoB, PsbK, RbcL2, RbcL1, and ITS2. *H. parviflora* was characterized by amplification of three DNA barcoding markers, namely, trnHpsbA1, PsbA-trnH2, and PsbK. The MatK1, RpoC, PsbK, RbcL2, RbcL1, and ITS2 markers of V. chinensis were amplified. V. macrocarpa was amplified by seven DNA barcoding markers: MatK1, RpoC, PsbZ-trnfM, trnHpsbA1, PsbK, PsbA-trnH2, and RbcL1. Finally, *D. bourdiollon* was amplified with ten DNA barcoding markers, generating barcodes with MatK1, RpoC, RpoB, RpoB-trnCGAR, trnHpsbA1, PsbK, RbcL2, ITS1, PsbA-trnH2, and RbcL1. The observed amplification and sequencing rates for each species were as follows: *V. indica* (30.76 ± 100), S. roxburghii (46.15 ± 100), *H. ponga* (69.23 ± 77.8), *D. indicus* (41.17 ± 100), *H. parviflora* (23.07 ± 100), *V. chinensis* (35.29 ± 100), *V. macrocarpa* (41.17 ± 100), and *D. bourdiollon* (70.58 ± 83.3). In the Ebenaceae family, three species were examined: *D. paniculata, D. crassiflora*, and *D. candolleana*. *D. paniculata* exhibited amplification with RpoC and RbcL2 DNA barcoding markers, while *D. crassiflora* showed amplification with eleven DNA barcoding markers, including RpoB, RpoB-trnCGAR, PsbZ-trnfM, trnHpsbA1, PsbK, RPB2, MatK2, PsbA-trnH2, RbcL1, Atp, and ITS2. D. candolleana, on the other hand, was amplified by the RpoC, trnHpsbA1, PsbK, RbcL2, and RbcL1 DNA barcoding markers (Table 3). The amplification and sequencing rates were 11.76 ± 100 for D. paniculata, 84.61 ± 100 for *D.* crassiflora and 29.41 ± 100 for D. candolleana. The Fabaceae family was represented by five species: *D. latifolia, P. marsupium, S. asoca, K. pinnatum*, and *P. santalinus*. *D. latifolia* was amplified and sequenced with nine DNA barcoding markers, namely, RpoB, RpoB-trnCGAR, PsbZ-trnfM, trnHpsbA1, RPB2, PsbA-trnH2, RbcL1, Atp and ITS2. Four DNA barcoding markers were used to amplify P. marsupium: RpoB, trnHpsbA1, PsbA-trnH2 and RbcL1. *S. asoca* was amplified with RpoB, trnHpsbA1, RPB2, PsbA-trnH2 and RbcL1. *K. pinnatum* was amplified and sequenced with RpoB, RpoB-trnCGAR, PsbA-trnH2, and RbcL1. Finally, *P. santalinus* was amplified with five barcoding markers: RpoB, TrnHpsbA1, PsbA-trnH2, RbcL1, and Atp (Table 3). The amplification and sequencing rates *were* 76.92± 90.8 *for D. latifolia,* 30.76± 100.0 *for P. marsupium,* 46.15± 83.3 *for S. asoca,* 61.53± 37.5 for K. pinnatum, and 38.46± 100% *for P. santalinus* (Fig). Within the Flacourtiaceae family, *H. macrocarpa* was the focal species subjected to amplification and sequencing utilizing two markers, RpoC and RbcL1, with an amplification and sequencing rate of 11.76 ± 100%. In the Lauraceae family, *C. riparium* emerged as the prominent species, displaying the highest amplification success in this investigation. Among the sixteen markers analyzed, fourteen were effectively amplified. These markers included MatK1, RpoC, RpoB, PsbZ-trnfM, trnHpsbA1, PsbK, RbcL2, PRK, RPB2, MatK2, PsbA-trnH2, RbcL1, Atp, and ITS2, with an amplification and sequencing rate of 88.23 ± 86.7%. *E. ribes* represents the Primulaceae family and was successfully amplified by twelve markers: MatK1, RpoC, RpoB, RpoB-trnCGAR, PsbZ-trnfM, trnHpsbA1, PsbK, RbcL2, PsbA-trnH2, RbcL1, Atp, and ITS2. The amplification and sequencing rates for E. ribes were 70.58 ± 91.7%. Within the Malvaceae family, three species have been identified: *P. reticulatum, S. occidentale* and *S. travancoricum*. Three DNA barcoding markers, RpoB, PsbK, and RbcL1, were successfully amplified from P. reticulatum. On the other hand, S. occidentale displayed amplification and sequencing of thirteen markers, namely, MatK1, RpoC, RpoB, RpoB-trnCGAR, PsbZ-trnfM, trnHpsbA1, PsbK, RbcL2, PRK, RPB2, PsbA-trnH2, RbcL1, and Atp. Similarly, three DNA barcoding markers were used to amplify and sequence S. travancoricum: RpoB, PsbZ-trnfM, and RbcL1. The amplification and sequencing rates for each species were recorded as follows: *P. reticulatum* (76.92 ± 100), *S. occidentale* (70.58 ± 100), and *S. travancoricum* (46.15 ± 57.1). In this study, *D. malabaricum*, which is a member of the Meliaceae family, was effectively amplified through four markers: MatK1, PsbK, RbcL1, and Atp. The recorded rate of amplification and sequencing for *D. malabaricum* was 46.15 ± 57.1%. Four species from the Myristicaceae family were examined in this study: *G. canarica, M. malabarica, M. beddomei* and *K. attenuata*. Commonly amplified markers across all species included trnHpsbA1, PsbA-trnH2, PsbZ-trnfM, and PsbK. The seven barcode markers of G. canarica were amplified and sequenced: MatK1, RpoC, RpoB, and PsbA-trnH2. *M. malabarica* was amplified by RpoB, RpoB-trnCGAR, RPB2, MatK2, RbcL1, and Atp. *M. beddomei* exhibited amplification by RpoC, RbcL2, and Atp. K. attenuata showed amplification of RbcL1 and Atp. The observed amplification and sequencing rates were as follows: *G. canarica* (52.94 ± 77.8), *M. malabarica* (76.92 ± 100), *M. beddomei* (52.94 ± 77.8) and *K. attenuata* (69.23 ± 66.7). In the Sapotaceae family, two species were examined: *M. bourdillonii* and *M. insignis*. M. bourdillonii was subjected to amplification and sequencing using the RpoC, PsbZ-trnfM, trnHpsbA1, PsbK, RbcL2, PsbA-trnH2, and Atp primers. *M. insignis* was amplified with RpoB, RpoB-trnCGAR, PsbZ-trnfM, trnHpsbA1, RPB2, PsbA-trnH2, RbcL1, and Atp. The amplification and sequencing rates were 52.94 ± 88.9 for *M.* bourdillonii and 30.76 ± 88.9 for M. insignis (Fig.; Table).

**Fig. 4.**
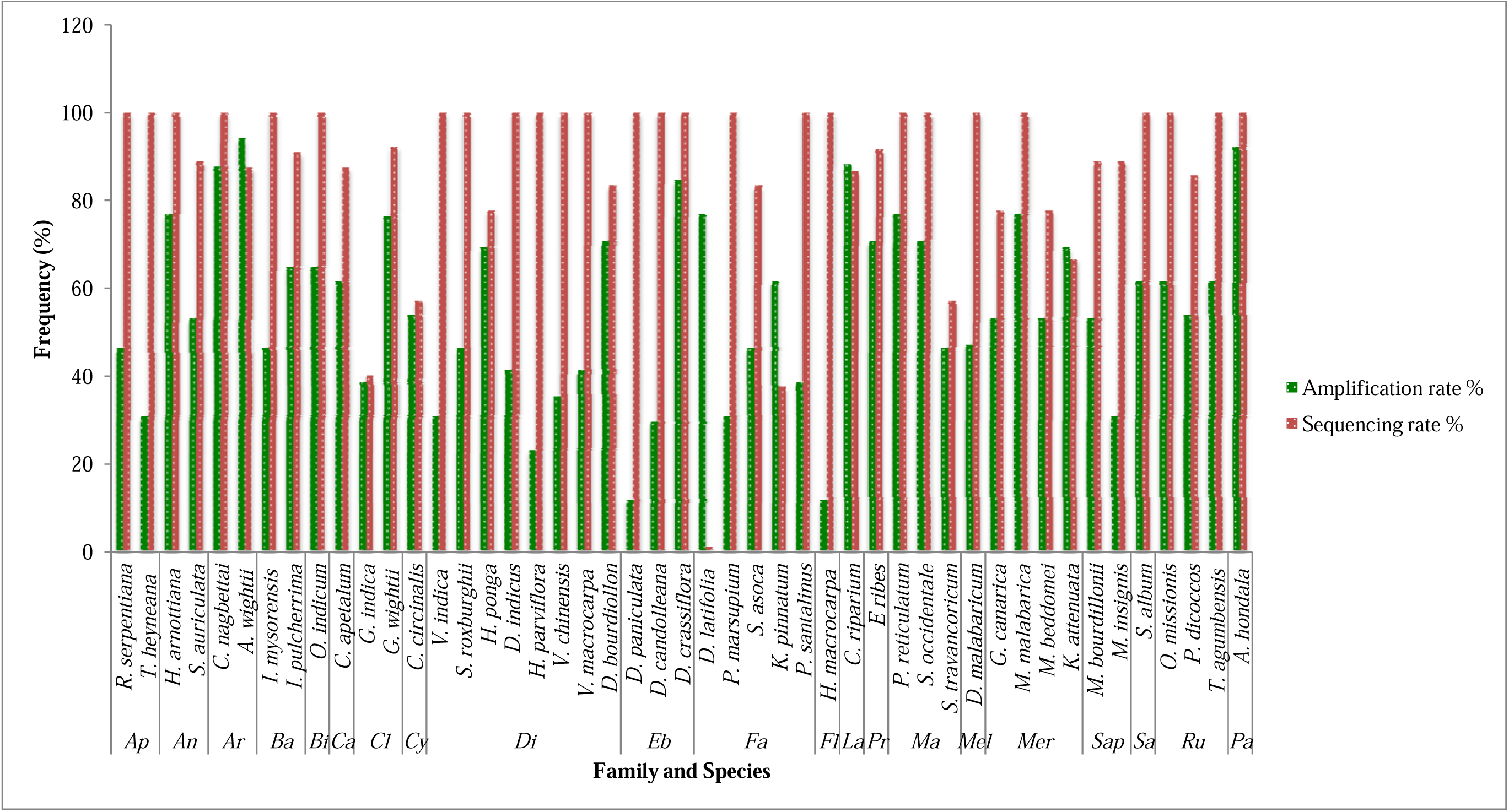
The successs rate of PCR amplification and sequencing of the sixteen barcode fragments across the 47 species from 21 families.

**Table 3.**
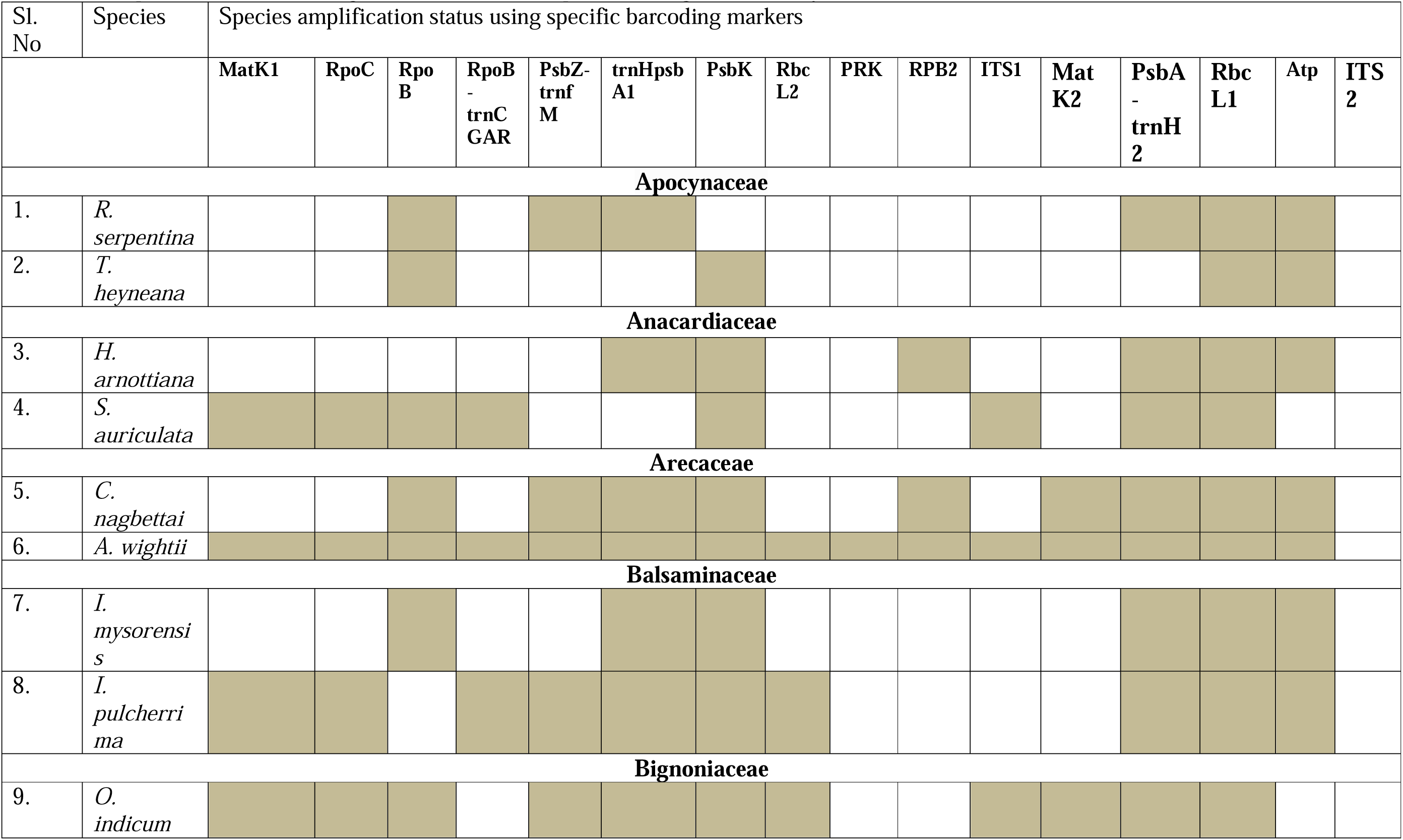
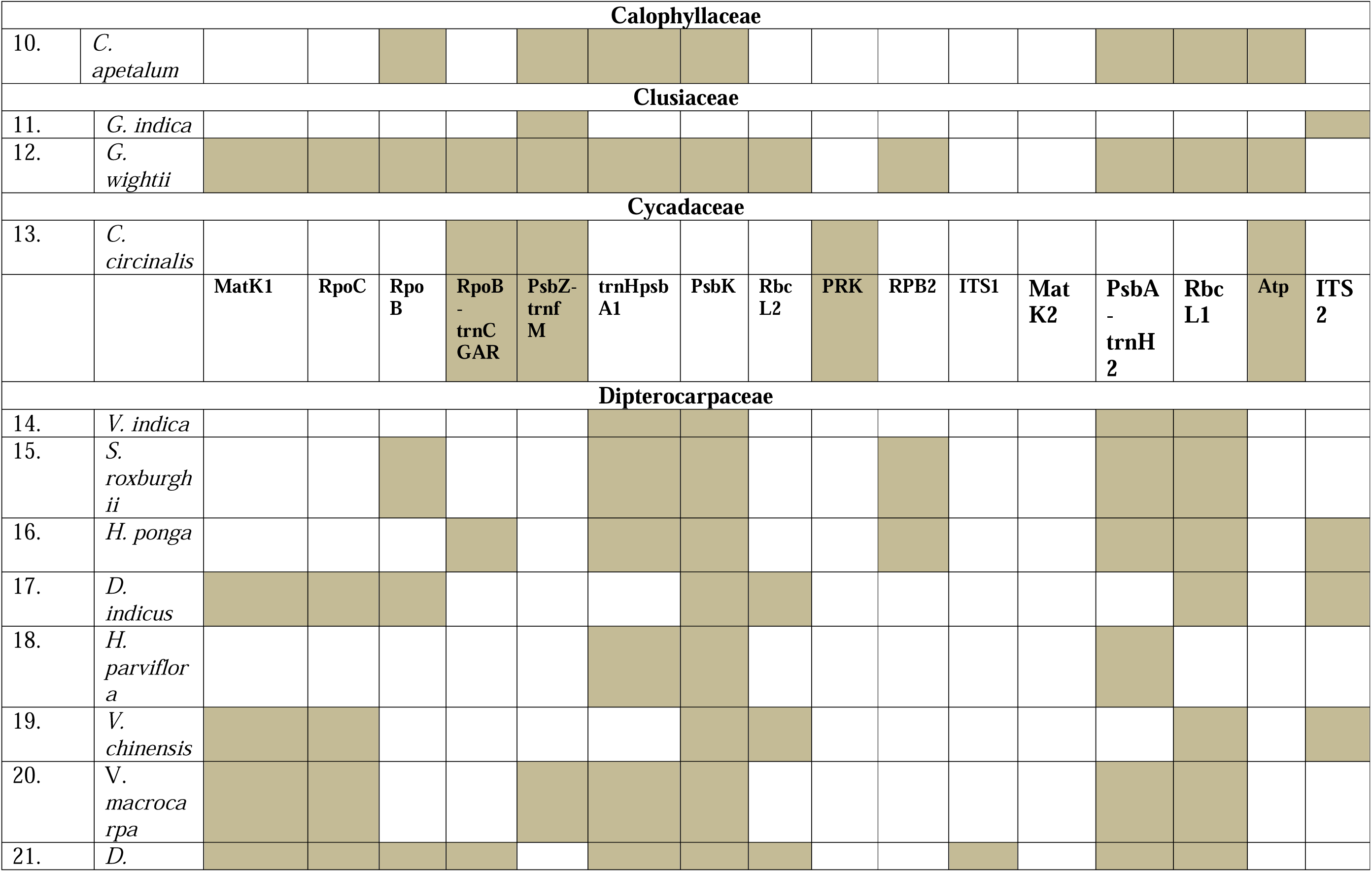
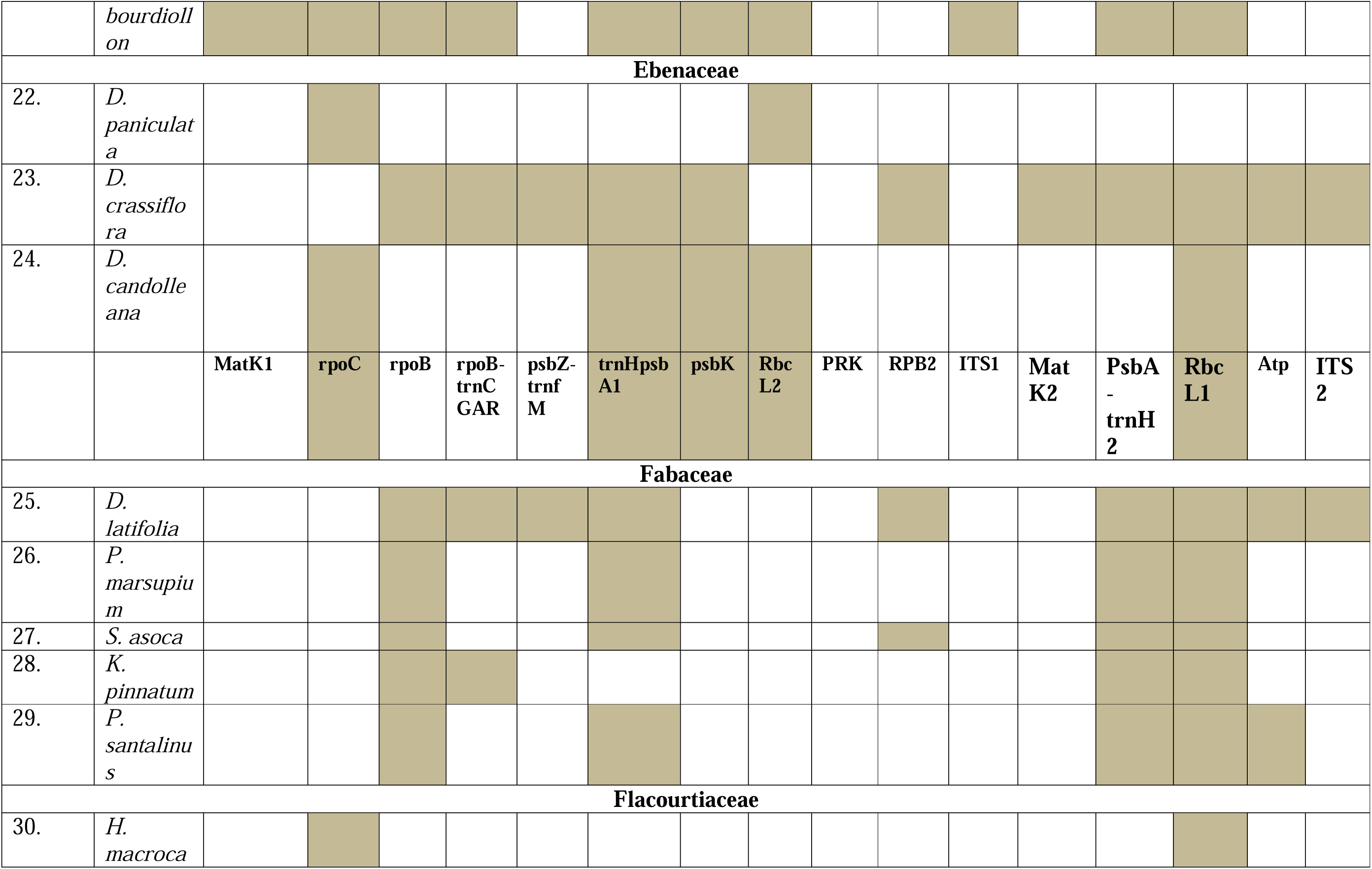
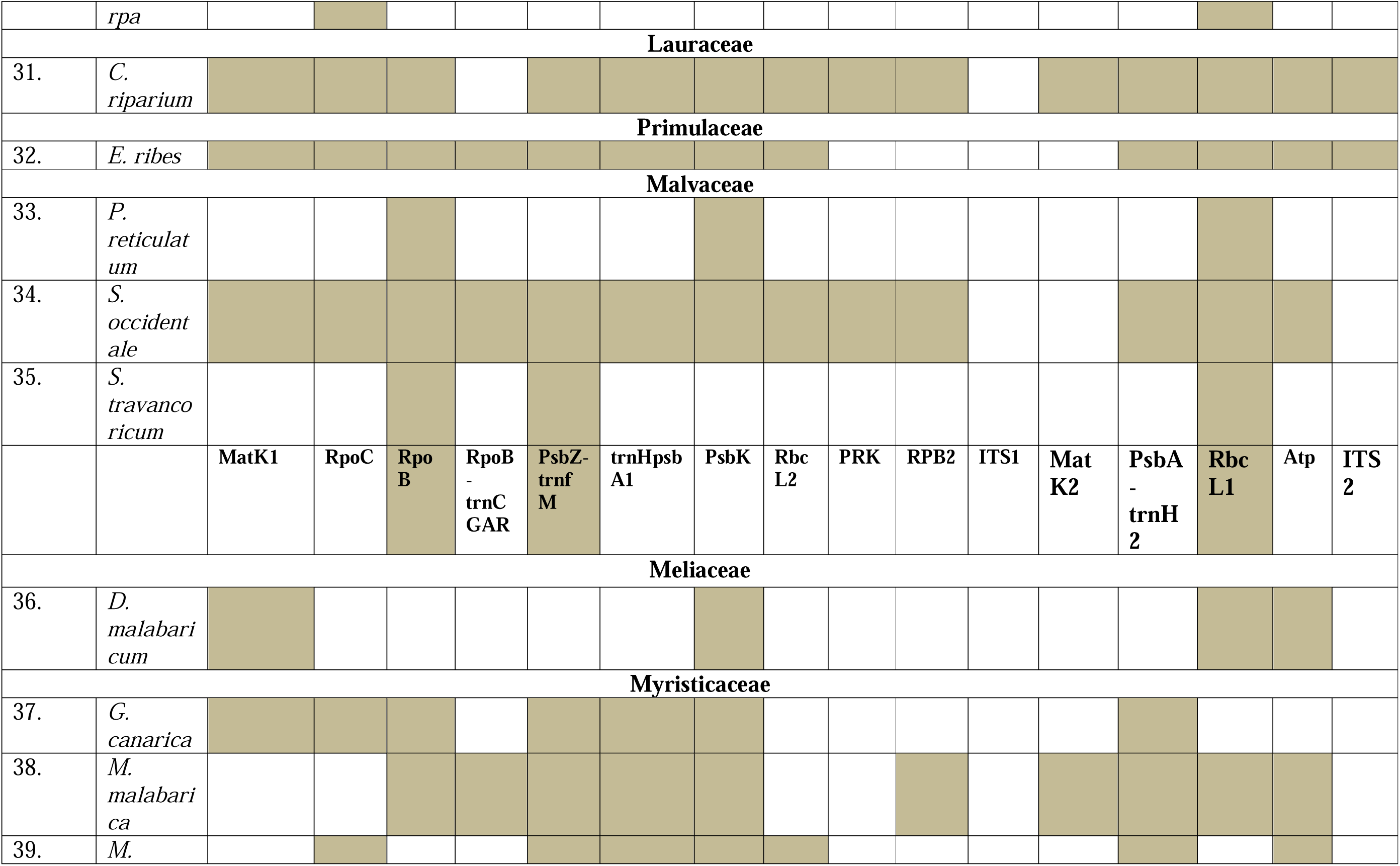
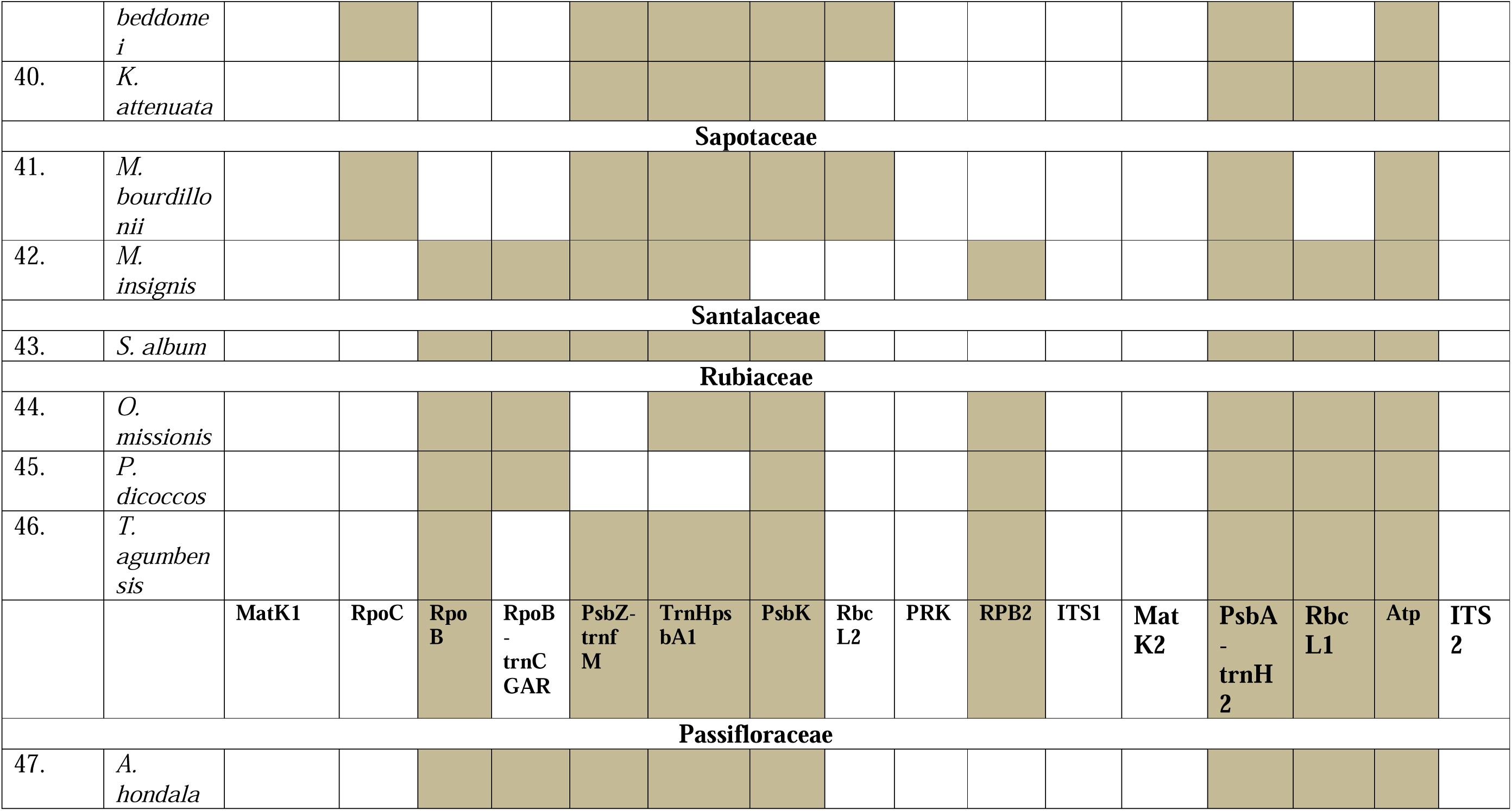
Amplified DNA Barcoding markers in selected species belongs to 21 family.

Within the Santalaceae family, the species *S. album* was included in this investigation. The gene was amplified using eight DNA barcoding markers: RpoB, RpoB-trnCGAR, PsbZ-trnfM, trnHpsbA1, PsbK, PsbA-trnH2, RbcL1, and Atp. The observed amplification rate for *S. album* was 61.53 ± 100%. In the Rubiaceae family, three species were examined: *O. missionis, P. dicoccos,* and *T. agumbensis*. These genes shared commonly amplified markers, including RpoB, PsbK, RPB2, PsbA-trnH2, RbcL1, and Atp. Additionally, RpoB-trnCGAR was amplified in *O. missionis* and *P. dicoccos*, while trnHpsbA1 was amplified in *O. missionis* and *T. agumbensis*. PsbZ-trnfM was specifically amplified in *T. agumbensis*. The amplification and sequencing rates were as follows: *O. missionis* (61.53 ± 100), *P. dicoccos* (53.84 ± 85.7), and *T. agumbensis* (61.53 ± 100). In this study, *A. hondala* was representative of the Passifloraceae family. The species were amplified and sequenced using eight DNA barcoding markers: RpoB, RpoB-trnCGAR, PsbZ-trnfM, trnHpsbA1, PsbK, PsbA-trnH2, RbcL1, and Atp. The amplification and sequencing rates were 92.34 ± 100 (Fig. 4; Table 3).

### Molecular analysis of species identification using DNA barcodes

Among the 47 species examined, 28 lacked DNA barcodes or any associated sequences, as outlined in Table 4. In this study, species without existing DNA barcodes underwent amplification and sequencing utilizing selected DNA barcoding markers, resulting in a diverse array of sequences. These sequences were subsequently compiled and collectively subjected to BLAST analysis to identify the top hits specific to each species. By constraining the database and refining it based on a pairwise identity (PI) threshold of more than 95%, several significant matches were identified. For instance, *T. heyneana* exhibited a high similarity with *Tabernaemontana bovina* (NC079611.1; PI=98.86%). Similarly, *H. arnottiana* and *S. auriculata*, which belong to the same family, were matched with *Semecarpus reticulatus* (NC069617.1; PI=98.82% and 98.94%, respectively). *A. wightii* was matched with *Wallichia disticha* (ON248714.1; PI=97.72%). Additionally, *I. mysorensis* and *I. pulcherrima*, from the same family, were matched with *I. hawkeri* (NC048520.1; PI=98.02%) and *D. mespiliformis* (MZ274088.1; PI=98.73%), respectively. Furthermore, *C. apetalum* was matched with *Calophyllum soulattri* (NC_068749.1; PI=96.13%), and *G. wightii* was matched with *Kingiodendron pinnatum* (EU361987.1; PI=98.96%). In the Dipterocarpaceae family, *V. indica, H. ponga, D. indicus*, and *D. bourdiollon* (with PIs ranging from 95.36% to 98.50%) were matched with *Vatica bantamensis, Hopea reticulata, Holigarna arnottiana,* and *Vatica bantamensis* (NC_071231.1; NC_052744.1; MZ648043.1; NC071231.1), respectively. Moreover, *D. paniculata* and *D. candolleana* from the Ebenaceae family were matched with *Diospyros mespiliformis* and *Dipterocarpus littoralis* (MZ274088.1; PI=99.17% and NC_081465.1; 99.49%). *K. pinnatum* was matched with *P. oxyphylla* (MZ274107.1; PI=95.63%), and *H. macrocarpa* was matched with *H. hainanensis* (NC042720.1; PI=98.98%). Furthermore, *C. riparium* was matched with *Cinnamomum verum* (MG280937.1; PI=99.54%). In the Malvaceae family, *P. reticulatum, S. occidentale*, and *S. travancoricum* were matched with *Pterospermum heterophyllum, Semecarpus australiensis*, and *Syzygium grijsii* (ON100918.1; PI=98.83%, AY594479.1; 97.74%, and NC_065156.1; 99.23%, respectively). Additionally, *D. malabaricum* was matched with *Tabernaemontana divaricata* (MZ_07333.1; PI=99.07%), and *G. canarica* was matched with *M. teysmannii* (NC_079584.1; PI=100%). Furthermore, *M. bourdillonii* and *M. insignis* from the Sapotaceae family were matched with *Impatiens hawkeri* and *Madhuca hainanensis* (NC_048520.1; PI=98.64% and NC_053619.1; PI=99.23%, respectively). *O. missionis, P. dicoccos*, and *T. agumbensis* were matched with *Atalantia ceylanica, Psydrax obovata,* and *P. rubra* (NC_065396.1; PI=99.77%, KY378666.1; 98.66%, and MZ958829.1; 99.85%, respectively). Finally, *A. hondala* was matched with *Adenia mannii* (KF724302.1; PI=98.97%) (Table 4; Supplementary file 1). The obtained nucleotide sequences were submitted to the DNA databank (https://www.ncbi.nlm.nih.gov/guide/dna-rna/) (Supplementary file 2).

**Table 4.** BLAST results of the amplified sequences of 28 species of western ghats.

| SI No. | Species | Barcoding markers | PI% | Species matching | Accession number |
| --- | --- | --- | --- | --- | --- |
| **Family** |  |  |  |  |  |
| **Apocynaceae** |  |  |  |  |  |
| 1. | *T. heyneana* | RbcL1+ Atp+ PsbK+ RpoB | 98.86 | *Tabernaemontana bovina* | NC079611.1 |
| **Anacardiaceae** |  |  |  |  |  |
| 2. | *H. arnottiana* | RbcL1+ psbA_trnH1+ Atp+ PsbK+<br>psbZ_trnfM+ RPB2+ RpoB+ trnHpsbA2+<br>RpoB_trnCGAR+ ITS2 | 98.82 | *Semecarpus reticulatus* | NC069617.1 |
| 3. | *S. auriculata* | MatK2+ RpoC+ RbcL1+ psbA-trnH1+<br>ITS2+ PsbK+ RpoB+ RpoB-trnCGAR | 98.94 | *S. reticulatus* | NC069617.1 |
| **Arecaceae** |  |  |  |  |  |
| 4. | *A. wightii* | MatK2+ RbcL1+ MatK2+ psbAtrnH2+<br>Atp+ PsbK+ psbZtrnfM+ RPB2+ RpoB+<br>RpoC+ trnHpsbA1+ RbcL2+<br>RpoBtrnCGAR | 97.72 | *Wallichia disticha* | ON248714.1 |
| **Balsaminaceae** |  |  |  |  |  |
| 5. | *I. mysorensis* | RbcL1+ trnHpsbA1+ PsbK+ RpoB+ psbA-<br>trnH1 | 98.02 | *I. hawkeri* | NC048520.1 |
| 6. | *I. pulcherrima* | MatK2+ RbcL1+ trnHpsbA1+ Atp+ PsbK+<br>psbZtrnfM + RpoC+ RpoBtrnCGAR+<br>psbAtrnH2+ RbcL2 | 98.73 | *D. mespiliformis*<br>(*ebenaceae*) | MZ274088.1 |
| **Calophyllaceae** |  |  |  |  |  |
| 7. | *C. apetalum* | RbcL1+ Atp+ PsbK+ trnHpsbA1+ RpoB+<br>psbAtrnH2 | 96.13 | *Calophyllum soulattri* | NC_068749.1 |
| **Clusiaceae** |  |  |  |  |  |
| 8. | *G. wightii* | MatK2+ RbcL1+ psbAtrnH2+ trnHpsbA1+<br>RpoB+ RpoC+ RpoBtrnCGAR+ Atp+<br>PsbK+ RPB2+ RbcL2 | 98.96 | *Kingiodendron pinnatum* | EU361987.1 |

| **Dipterocarpaceae** |  |  |  |  |  |
| --- | --- | --- | --- | --- | --- |
| 9. | *V. indica* | RbcL1+ PsbA-trnH2+ trnHpsbA1+ PsbK | 98.50 | *Vatica bantamensis* | NC_071231.1 |
| 10. | *H. ponga* | RPB2+ trnHpsbA1+ RpoB_trnCGAR+ ITS2 | 96.97 | *Hopea reticulata* | NC_052744.1 |
| 11. | *D. indicus* | MatK2+ RbcL1+ PsbK+ RpoB+ RpoC+ ITS2+ RbcL2 | 98.48 | *Holigarna arnottiana* | MZ648043.1 |
| 12. | *D. bourdiollon* | MatK2+ RbcL1+ trnHpsbA1+ PsbK+ RpoB+ RpoC RpoB-trnCGAR+ psbAtrnH2+ RbcL2 | 95.36 | *Vatica bantamensis*) | NC071231.1 |
| **Ebenaceae** |  |  |  |  |  |
| 13. | *D. paniculata* | RpoC+ RbcL1 | 99.17 | *Diospyros mespiliformis* | MZ274088.1 |
| 14. | *D. candolleana* | RbcL1+ psbAtrnH2+ + PsbK+ RpoC+ RbcL2 | 99.49 | *Dipterocarpus littoralis* | NC_081465.1 |
| **Fabaceae** |  |  |  |  |  |
| 15. | *K. pinnatum* | trnHpsbA1+ RpoB+ RpoB_trnCGAR | 95.63 | *Prioria oxyphylla* | MZ274107.1 |
| **Flacourtiaceae** |  |  |  |  |  |
| 16. | *H. macrocarpa* | RbcL1+ RpoC | 98.98 | *H. hainanensis* | NC042720.1 |
| **Lauraceae** |  |  |  |  |  |
| 17. | *C. riparium* | MatK2+ RbcL1+ Atp+ PsbK+ RpoB+ RpoC+ RPB2+ PsbZ-trnfM+ trnHpsbA1+ ITS2 | 99.54 | *Cinnamomum verum* | MG280937.1 |
| **Malvaceae** |  |  |  |  |  |
| 18. | *P. reticulatum* | RbcL1+ PsbK+ RpoB | 98.83 | *Pterospermum heterophyllum* | ON100918.1 |
| 19. | *S. occidentale* | MatK2+ RbcL1+ psbAtrnH1 + Atp+ PsbK+ PsbZ-trnfM+ RpoB+ RpoC+ RpoB-trnCGAR+ RPB2+ psbAtrnH2+ RbcL2 | 97.74 | *Semecarpus australiensis* | AY594479.1 |
| 20. | *S. travancoricum* | RbcL1+ PsbZ-trnfM + RpoB+ psbAtrnH1 | 99.23 | *Syzygium grijsii* | NC_065156.1 |
| **Meliaceae** |  |  |  |  |  |
| 21. | *D. malabaricum* | MatK2+ Atp+ RpoC+ RbcL1+ PsbK+ RpoB+ rbcL2+ ITS2 | 99.07 | *Tabernaemontana divaricata* | MZ_07333.1 |

| **Myristicaceae** |  |  |  |  |  |
| --- | --- | --- | --- | --- | --- |
| **22.** | *G. canarica* | MatK2+ trnHpsbA1+ PsbK+ PsbZ-trnfM+ RpoC+ trnHpsbA_2+ RpoB | 100 | *M. teysmannii* | NC_079584.1 |
| **Sapotaceae** |  |  |  |  |  |
| **23.** | *M. bourdillonii* | RbcL1+ trnHpsbA1+ RpoC+ trnHpsbA_2+ Atp+ PsbK+ rbcL2 | 98.64 | *Impatiens hawkeri* | NC_048520.1 |
| **24.** | *M. insignis* | RbcL1+ trnHpsbA1+ Atp+ PsbZ-trnfM+ RpoB+ RpoB-trnCGAR + trnHpsbA_2+ RPB2 | 99.23 | *M. hainanensis* | NC_053619.1 |
| **Rubiaceae** |  |  |  |  |  |
| **25.** | *O. missionis* | Cn_RbcL+ Atp+ PsbK+ RpoB+ RpoB-trnCGAR+ trnHpsbA1+ RPB2 | 99.77 | *Atalantia ceylanica* | NC_065396.1 |
| **26.** | *P. dicoccos* | rbcL1+ Atp+ PsbK+ RpoB+ trnHpsbA1+ PsbA-trnH2 | 98.66 | *Psydrax obovata* | KY378666.1 |
| **27.** | *T. agumbensis* | Cn_RbcL+ Atp+ PsbA-trnH2+ PsbK+ RPB2+ RpoB+ PsbZ-trnfM | 99.85 | *Psychotria rubra* | MZ958829.1 |
| **Passifloraceae** |  |  |  |  |  |
| **28.** | *A. hondala* | RbcL1+ trnHpsbA1+ Atp+ psbZ-trnfM+ RpoB+ RpoB-trnCGAR+ trnHpsbA2 | 98.97 | *Adenia mannii* | KF724302.1 |

## Discussion

Our findings indicated that the dissimilarities between species become more disparate when more independent loci are used. In this study, our aim was to generate DNA barcode sequences for endangered, threatened, and vulnerable forest species that are challenging to identify morphologically. The success rate of this study shows that barcoding markers can accurately distinguish between species within a family. In our study, the highest amplification rates were detected for rbcL1 (40 species), psbtrnH2 (36 species), and PsbA-trnH1 (33 species) (Table 3). These findings suggested that these markers could serve as promising universal identifiers for the majority of the selected forest species. Similar results were reported by Kang et al. (2017) and Huang et al. (2015), indicating the potential universality of the *rbcL and* PsbA-trnH1 barcodes. However, the majority lacked available DNA barcodes. Furthermore, in cases where DNA barcodes were available, the number of sequences was limited. In our study, we expanded upon existing DNA barcodes by developing additional markers for species with available sequences. Additionally, we successfully generated DNA barcode sequences for the majority of endangered species in the Western Ghats region for which barcode sequences were previously unavailable. For the 19 species with existing DNA barcodes and sequences, we expanded the available data by generating additional DNA sequences using a broader array of DNA barcoding markers. According to the literature, some DNA barcodes were accessible for the following species: viz; *R. serpentina*; trnL-trnF, ITS2, matK, rbcL and trnL (Eurlings et al. (2013); Cahyaningsih, et al. (2022); *C. nagbettai*; rbcL, matK, psbA-trnH, rpoC, rpoB, psbKpsbI, atpF-atpH and RPB2 (Kurian, et al. (2020); (Table 1); *O. indicum*; ITS2, psbA-trnH (Cui, et al. (2020); *G. indica*; rbcL, ITS, psbA-trnHf, RpoB-trnCGAR, Matk (Seethapathy et al. 2018); C. *circinalis*; ITS2 (Gao, et al. (2010); *S. roxburghii*; ITS1 and ITS2; psbM, rbcL, psbH, rpoC2 (Osathanunkul, et al. (2021); *H. parviflora;* MatK, rbcL (data unpublished); *V. chinensis;* rbc However, in our study, we developed additional markers beyond those that currently exist. We observed that certain existing markers were not amplified in our samples, possibly due to sequence mixing, as evidenced in species such as *R. serpentina, G. indica, C. circinalis, S. roxburghii*, and *P. santalinus* (Table 1). Furthermore, our study revealed lower amplification results with the ITS marker compared to findings reported in the literature (Selvaraj et al. 2014). Our efforts resulted in successful DNA barcoding for 28 species previously lacking representation in the database. The accuracy of the DNA barcodes was determined by comparing the molecular identification results obtained via BLAST searches against available reference databases. The number of matched genes was greatest at the genus level for all amplified and sequenced markers (Table 4; Supplementary file 1). The analysis revealed that the most precise level of identification closely resembled that of their respective neighboring species within the same family across all the amplified and sequenced markers (Supplementary file 1). However, while some species were confidently matched at the genus level, others could only be identified up to the family level. This discrepancy might stem from variations in genetic markers or the availability of reference sequences. Conversely, when individually analyzing the sequences, specific markers such as *A. wightii, V. indica*, and *D. bourdiollon* provided genus-level matches. Additionally, some species, including *G. wightii, D. indicus, D. malabaricum* and *M. bourdillonii*, unexpectedly showed matches with different families (Supplementary file 1).

## Conclusion

The findings of this study contribute to the establishment of a database encompassing threatened forest species not only in western Ghats but also in other regions. This study can improve forest biodiversity monitoring by detecting species at the molecular level. Such a database can play a pivotal role in enhancing conservation endeavors by facilitating the precise identification and monitoring of endangered species of western Ghats. In our research, the most significant amplification rates were noted for rbcL1 (40 species), psbtrnH2 (36 species), and PsbA-trnH1 (33 species). DNA amplification ranged from 11.76% to 94.11%. This highlights the ongoing need to expand DNA barcoding initiatives to encompass a wider array of species, thereby improving species identification accuracy and supporting conservation endeavors.

## Acknowledgments

The authors express gratitude to the Director General, Director Research, Head of the Centre for Forestry Ecology and Wildlife at the Environmental Management and Policy Research Institute. Furthermore, the authors gratefully acknowledge the Forest Department of Karnataka and the forest officers for generously granting permission to collect samples from the protected region and for their valuable encouragement throughout this study.

## Data availability statement

The authors declare that all the data supporting the findings of this study are available from GenBank (https://www.ncbi.nlm.nih.gov/genbank/).

