## Supplementary files for "DNA barcode developement based on chloroplast and ITS genes for species identification of endangered and threated species of Western Ghats, India"

Supplementary File 1. Barcodes not available (Supplementary file)

| **Sl No.** | **Species** | **Barcoding markers** | **PI%** | **Species matching** | **Accession number** |
|---|---|---|---|---|---|
| **Family** | | | | | |
| **(Apocynaceae)** | | | | | |
| **1.** | *T. heyneana* | RbcL1 | 99.31 | *Tabernaemontana divaricate* | X91772.1 |
| | | Atp | 98.40 | *T. divaricate* | MN044037.1 |
| | | PsbK | 97.27 | *T. divaricate* | MZ073339.1 |
| | | RpoB | 99.15 | *T. bovina* | NC079611.1 |
| | | Overall matching | 98.86 | *Tabernaemontana bovina* | NC079611.1 |
| **Anacardiaceae** | | | | | |
| **2.** | *H. arnottiana* | RbcL1 | 99.34 | *Mangifera indica* | MN711724.1 |
| | | PsbAtrnH1 | 96.48 | *Semecarpus sp.* | LC737830.1 |
| | | Atp | 94.87 | *Semecarpus reticulatus* | NC_069617.1 |
| | | PsbK | 94.87 | *S. reticulatus* | NC_069617.1 |
| | | Cn_psbZ_trnfM | 96.87 | *S. reticulatus* | NC_069617.1 |
| | | RPB2 | 98 | *Macrosolen cochinchinensis* | KP263343.1 |
| | | RpoB | 99.78 | *S. reticulatus* | NC069617.1 |
| | | Ci_trnHpsbA | 96.31 | *S. reticulatus* | NC069617.1 |
| | | Ga_RpoB_trnCGAR | 99.40 | *S. reticulatus* | NC069617.1 |
| | | ITS2 | 97.50 | *Dalbergia latifolia* | MH465107.1 |
| | | Overall matching | 98.82 | *Semecarpus reticulatus* | NC069617.1 |
| **3.** | *S. auriculata* | Matk_1 | 99.85 | *Holigarna arnottiana* | MZ648043.1 |
| | | RpoC | 99.37 | *Semecarpus reticulatus* | NC069617.1 |
| | | RbcL1 | 99.14 | *Melanochyla fulvinervia* | MH332443.1 |
| | | PsbA-trnH1 | 97.70 | *Holigarna beddomei* | OP476725.1 |
| | | ITS2 | 98.67 | *H. beddomei* | OP474007.1 |
| | | PsbK | 96.14 | *S. reticulatus* | NC069617.1 |
| | | RpoB | 99.78 | *S. reticulatus* | NC069617.1 |
| | | RpoB-trnCGAR | 98.94 | *S. reticulatus* | NC069617.1 |
| | | overall | 98.94 | *Semecarpus reticulatus* | NC069617.1 |

| | | | | | |
|---|---|---|---|---|---|
| **Arecaceae** | | | | | |
| **4.** | *A. wightii* | Matk_1 | 99.38 | *Arenga undulatifolia* | AM114593.1 |
| | | RbcL1 | 99.66 | *A. pinnata* | AY012478.1 |
| | | Matk_1 | 99.76 | *A. hookeriana* | AM114592.1 |
| | | PsbAtrnH_2 | 99.56 | *A. wightii* | JF345043.1 |
| | | Atp | 98.14 | *A. westerhoutii* | NC_079705.1 |
| | | PsbK | 98.80 | *A. pinnata* | NC045907.1 |
| | | PsbZ-trnfM | 99.53 | *Wallichia gracilis* | NC079761.1 |
| | | RPB2 | 98.53 | *A. pinnata* | HQ720487.1 |
| | | RpoB | 99.08 | *A. micrantha* | MW044623.1 |
| | | RpoC | 99.36 | *Wallichia caryotoides* | NC_079759.1 |
| | | TrnHpsbA2 | 99.71 | *A. wightii* | JF345043.1 |
| | | RbcL_2 | 99.48 | *A. hookeriana* | MG437585.1 |
| | | RpoBtrnCGAR | 97.72 | *Wallichia disticha* | ON248714.1 |
| | | Overall matching | 97.72 | *Wallichia disticha* | ON248714.1 |
| **Balsaminaceae** | | | | | |
| **5.** | *I. mysorensis* | RbcL1 | 98.66 | *Impatiens gardneriana* | MW459331.1 |
| | | PsbA-trnH1 | 97.30 | *I. hawkeri* | NC_048520.1 |
| | | PsbK | 93.26 | *I. hawkeri* | NC_048520.1 |
| | | RpoB | 99.57 | *I. hawkeri* | NC_048520.1 |
| | | Cn_ psbA-trnH | 98.73 | *I. hawkeri* | NC_048520.1 |
| | | Overall | 98.02 | *I. hawkeri* | NC048520.1 |
| **6.** | *I. pulcherrima* | Matk_1 | 99.39 | *Diospyros scalariformis* | EU980975.1 |
| | | RbcL1 | 98.33 | *Diospyros argentea* | GU471704.1 |
| | | PsbAtrnH1 | 96.39 | *Diospyros morrisiana* | KP095202.1 |
| | | Atp | 96.88 | *Diospyros virginiana* | NC_039555.1 |
| | | PsbK | 99.75 | *Heritiera elata* | NC_043925.1 |
| | | PsbZ-trnfM | 99.60 | *Diospyros oleifera* | NC_030787.1 |
| | | RpoC | 99.79 | *Diospyros mespiliformis* | MZ274088.1 |
| | | RpoBtrnCGAR | 98.96 | *Heritiera elata* | NC_043925.1 |
| | | trnHpsbA2 | 95.14 | *Diospyros glaucifolia* | HQ427082.1 |

| | | | | | |
|---|---|---|---|---|---|
| | | RbcL2 | 98.10 | *Diospyros conocarpa* | MN366481.1 |
| | | Overall | 98.73 | *D. mespiliformis* | MZ274088.1 |
| | | | **Calophyllaceae** | | |
| **7.** | *C. apetalum* | RbcL1 | 99.49 | *Calophyllum inophyllum* | KX807131.1 |
| | | Atp | 97.60 | *C. inophyllum* | NC_067766.1 |
| | | PsbK | 98.91 | *C. inophyllum* | - |
| | | PsbAtrnH1 | 95.20 | *Calophyllum polyanthum* | MW047955.1 |
| | | RpoB | 99.12 | *Calophyllum soulattri* | NC_068749.1 |
| | | **psbZ-trnfM** | 96.13 | *C. soulattri* | - |
| | | trnHpsbA2 | 96.53 | *C. soulattri* | - |
| | | overall | 96.13 | *Calophyllum soulattri* | - |
| | | | **Clusiaceae** | | |
| **8** | *G. wightii* | Matk_1 | 98.96 | *Kingiodendron pinnatum* | EU361987.1 |
| | | RbcL1 | 99.23 | *K. novoguineense* | JF739130.1 |
| | | PsbAtrnH2 | 91.57 | *Brandzeia filicifolia* | MN992879.1 |
| | | trnHpsbA1 | 90.37 | *B. filicifolia* | - |
| | | PsbZ-trnfM | 99.31 | *Wallichia gracilis* | NC_079761.1 |
| | | RpoB | 99.34 | *Prioria oxyphylla* | MZ274107.1 |
| | | RpoC | 98.94 | *P. oxyphylla* | - |
| | | RpoBtrnCGAR | 96 | *P. oxyphylla* | - |
| | | Atp | 92.38 | *P. oxyphylla* | - |
| | | PsbK | 96.08 | *P. oxyphylla* | - |
| | | RPB2 | | NA | |
| | | RbcL2 | 98.43 | *Senegalia modesta* | MF694648.1 |
| | | Overall | 98.96 | *Kingiodendron pinnatum* | EU361987.1 |
| | | | **Dipterocarpaceae** | | |
| **9.** | *V. indica* | RbcL1 | 99.48 | *Vateria copallifera* | KY973259.1 |
| | | PsbA-trnH2 | 98.58 | *V. copallifera* | KR338463.1 |
| | | trnHpsbA1 | 97.66 | *V. copallifera* | - |
| | | PsbK | 98.63 | *Vatica mangachapoi* | NC_041485.1 |
| | | overall | 98.50 | *Vatica bantamensis* | NC_071231.1 |

| | | | | | |
|---|---|---|---|---|---|
| **10.** | *H. ponga* | RPB2 | 94.34 | *Artabotrys zeylanicus* | MN231406.1 |
| | | trnHpsbA2 | 99.62 | *Hopea chinensis* | MW074190.1 |
| | | RpoB_trnCGAR | 96.97 | *Hopea reticulata* | NC_052744.1 |
| | | ITS2 | 98.67 | *Dalbergia latifolia* | MH465107.1 |
| | | overall | 96.97 | *Hopea reticulata* | NC_052744.1 |
| **11.** | *D. indicus* | Matk1 | 98.48 | *Holigarna arnottiana* | MZ648043.1 |
| | | RbcL1 | 97.57 | *Mangifera indica* | MW164971.1 |
| | | PsbK | 96.22 | *Semecarpus reticulatus* | NC_069617.1 |
| | | RpoB | 97.86 | *S. reticulatus* | - |
| | | RpoC | 99.58 | *S. reticulatus* | - |
| | | ITS2 | 97.94 | *Holigarna beddomei* | OP474007.1 |
| | | rbcL2 | 98.53 | *Humulus lupulus* | MF973024.1 |
| | | overall | 98.48 | *Holigarna arnottiana* | MZ648043.1 |
| **12.** | *D. bourdiollon* | Matk1 | 99.86 | *Vateria copallifera* | KY973069.1 |
| | | RbcL1 | 99.12 | *Vatica odorata* | KY973280.1 |
| | | PsbA-trnH1 | 98.41 | *Vatica diospyroides* | MW074178.1 |
| | | PsbK | 94.72 | *Neobalanocarpus heimii* | EU918763.1 |
| | | RpoB | 98.25 | *Dipterocarpus littoralis* | OQ831430.1 |
| | | RpoC | 98.48 | *D. littoralis* | - |
| | | RpoB-trnCGAR | 97.00 | *Vatica guangxiensis* | NC051531.1 |
| | | PsbA-trnH2 | 98.43 | *Vatica diospyroides* | MW074177.1 |
| | | RbcL2 | 97.04 | *Afzelia bella* | MN521397.1 |
| | | overall | 95.36 | *Vatica bantamensis* | NC071231.1 |
| **Ebenaceae** | | | | | |
| **13** | *D. paniculata* | RpoC | 99.17 | *Diospyros mespiliformis* | MZ274088.1 |
| | | RbcL1 | 99.50 | *Diospyros melanoxylon* | KX344717.1 |
| | | overall | 99.17 | *Diospyros mespiliformis* | MZ274088.1 |
| **14.** | *D. candolleana* | RbcL1 | 99.49 | *Dipterocarpus littoralis* | NC_081465.1 |
| | | PsbAtrnH2 | 98.91 | *Dipterocarpus zeylanicus* | KR338464.1 |
| | | PsbK | 98.35 | *Dipterocarpus retusus* | NC_067812.1 |

| | | | | | |
|---|---|---|---|---|---|
| | | RpoC | 98.35 | *D. retusus* | NC_067812.1 |
| | | RbcL2 | 96.68 | *Euphorbia prostrata* | MF694713.1 |
| | | Overall | 99.49 | *Dipterocarpus littoralis* | NC_081465.1 |
| **Fabaceae** | | | | | |
| **15.** | *K. pinnatum* | trnHpsbA2 | 93.36 | *Prioria balsamifera* | MZ274106.1 |
| | | RpoB | 99.36 | *Prioria oxyphylla* | MZ274107.1 |
| | | RpoB_trnCGAR | 95.63 | *P. oxyphylla* | - |
| | | overall | 95.63 | *Prioria oxyphylla* | - |
| **Flacourtiaceae** | | | | | |
| **16.** | *H. macrocarpa* | RbcL1 | 98.98 | *Hydnocarpus hainanensis* | MH667433.1 |
| | | RpoC | 99.36 | *H. hainanensis* | NC_042720.1 |
| | | overall | 98.98 | *H. hainanensis* | NC042720.1 |
| **Lauraceae** | | | | | |
| **17.** | *C. riparium* | Matk1 | 100 | *Cinnamomum verum* | MG280937.1 |
| | | RbcL1 | 98.97 | *Cinnamomum cappara-coronde* | MK243400.1 |
| | | Atp | 99.21 | *Cinnamomum pittosporoides* | NC_048978.1 |
| | | PsbK | 99.74 | *Cinnamomum camphora* | MN689735.1 |
| | | RpoB | 99.36 | *C. camphora* | MN689735.1 |
| | | RpoC | 98.95 | *Cinnamomum aromaticum* | OL943971.1 |
| | | RPB2 | 97.78 | *C. aromaticum* | MN698963.1 |
| | | psbZ-trnfM | 97.43 | *Cinnamomum yabunikkei* | NC_044864.1 |
| | | trnHpsbA2 | 95.79 | *Cinnamomum travancoricum* | MF547522.1 |
| | | ITS2 | NA | | |
| | | overall | 99.54 | *Cinnamomum verum* | MG280937.1 |
| **Malvaceae** | | | | | |
| **18.** | *P. reticulatum* | RbcL1 | 99.83 | *Pterospermum xylocarpum* | KF381124.1 |
| | | PsbK | 98.77 | *Pterospermum menglunense* | NC_057978.1 |
| | | RpoB | 99.57 | *P. menglunense* | NC_057978.1 |
| | | Overall | 98.83 | *Pterospermum heterophyllum* | ON100918.1 |
| **19.** | *S. occidentale* | Matk1 | 99.08 | *Holigarna arnottiana* | MZ648043.1 |
| | | RbcL1 | 98.79 | *Mangifera casturi* | LC602987.1 |

| | | | | | |
|---|---|---|---|---|---|
| | | PsbA-trnH1 | 99.76 | *Semecarpus* sp. | GU080317.1 |
| | | Atp | 90.34 | *Semecarpus reticulatus* | NC_069617.1 |
| | | PsbK | 96.68 | *S. reticulatus* | - |
| | | psbZ-trnfM | 97.39 | *S. reticulatus* | - |
| | | RpoB | 100 | *S. reticulatus* | - |
| | | RpoC | 99.57 | *S. reticulatus* | - |
| | | RpoB-trnCGAR | 96.89 | *S. reticulatus* | - |
| | | RPB2 | 98 | *Macrosolen cochinchinensis* | KP263343.1 |
| | | trnHpsbA2 | 99.52 | *Semecarpus* sp. | GU080317.1 |
| | | RbcL2 | 97.50 | *Ophiorrhiza* sp. | MH185932.1 |
| | | Overall | 97.74 | ***Semecarpus australiensis*** | AY594479.1 |
| **20.** | *S. travancoricum* | RbcL1 | 99.14 | *Syzygium cumini* | KF381145.1 |
| | | psbZ-trnfM | 99.36 | *Syzygium grijsii* | NC_065156.1 |
| | | RpoB | 99.53 | *Syzygium oleosum* | XM_056308039.1 |
| | | PsbA-trnH2 | 99.34 | *Syzygium cumini* | LC_461892.1 |
| | | overall | 99.23 | *Syzygium grijsii* | NC_065156.1 |
| | | | **Meliaceae** | | |
| **21.** | *D. malabaricum* | Matk1 | 99.07 | *Tabernaemontana sphaerocarpa* | GU_973968.1 |
| | | Atp | 98.02 | *Tabernaemontana divaricate* | MN044037.1 |
| | | RpoC | 99 | *T. divaricate* | MZ073339.1 |
| | | RbcL1 | 100 | *T. divaricate* | MK933718.1 |
| | | PsbK | 100 | *T. divaricate* | - |
| | | RpoB | 99.57 | *Tabernaemontana bovina* | NC_079611.1 |
| | | rbcL2 | 99.82 | *T. bovina* | - |
| | | ITS2 | 95.62 | *Tabernaemontana bufalina* | KC878558.1 |
| | | Overall | 99.07 | ***Tabernaemontana divaricata*** | MZ_07333.1 |
| | | | **Myristicaceae** | | |
| **22.** | *G. canarica* | Matk1 | 98.45 | *Virola michelii* | FJ038737.1 |
| | | PsbAtrnH2 | 98.03 | *Knema attenuata* | OK053228.1 |
| | | PsbK | 98.79 | *Myristica fatua* | OP866725.1 |
| | | psbZ-trnfM | 98.26 | *Knema tenuinervia* | NC_060714.1 |

| | | | | | |
|---|---|---|---|---|---|
| | | RpoC | 100 | *Knema tenuinervia* | - |
| | | trnHpsbA2 | 96.52 | *K. tenuinervia* | - |
| | | RpoB | 98.90 | *Myristica teysmannii* | NC_079584.1 |
| | | Overall | 100 | *M. teysmannii* | - |
| | | | **Sapotaceae** | | |
| **23.** | *M. bourdillonii* | RbcL1 | 97.41 | *Impatiens balsamina* | MZ902354.1 |
| | | PsbAtrnH_2 | 95.64 | *Impatiens hawkeri* | NC_048520.1 |
| | | RpoC | 98.98 | *I. hawkeri* | NC_048520.1 |
| | | trnHpsbA1 | 96.07 | *I. hawkeri* | NC_048520.1 |
| | | Atp | 95.03 | *Impatiens platysepala* | NC_068751.1 |
| | | PsbK | 96.47 | *Impatiens balsamina* | NC_059942.1 |
| | | rbcL_2 | 98.09 | *Impatiens piufanensis* | NC_037401.1 |
| | | overall | 98.64 | *Impatiens hawkeri* | NC_048520.1 |
| **24.** | *M. insignis* | RbcL1 | 99.32 | *Mimusops elengi* | MK947210.1 |
| | | PsbA-trnH1 | 99.62 | *Madhuca hainanensis* | AM179725.1 |
| | | Atp | 99.05 | *M. hainanensis* | NC_053619.1 |
| | | psbZ-trnfM | 97.64 | *M. hainanensis* | NC_053619.1 |
| | | RpoB | 99.09 | *M. hainanensis* | NC_053619.1 |
| | | RpoB-trnCGAR | 99.23 | *M. hainanensis* | NC_053619.1 |
| | | PsbA-trnH2 | 99.36 | *Palaquium microphyllum* | LC737417.1 |
| | | RPB2 | | NA | |
| | | overall | 99.23 | *M. hainanensis* | NC_053619.1 |
| | | | **Rubiaceae** | | |
| **25.** | *O. missionis* | RbcL1 | 98.68 | *Aegle marmelos* | OP102127.1 |
| | | Atp | 97.87 | *Atalantia buxifolia* | MZ408683.1 |
| | | PsbK | 98.17 | *Atalantia ceylanica* | NC_065396.1 |
| | | RpoB | 96.41 | *A. ceylanica* | NC_065396.1 |
| | | RpoB-trnCGAR | 96.77 | *A. ceylanica* | NC_065396.1 |
| | | TrnHpsbA2 | 85.85 | *Citrus limonia* | NC_065755.1 |
| | | RPB2 | 96 | *Citrus clementina* | XM_006423531.2 |
| | | overall | 99.77 | *Atalantia ceylanica* | NC_065396.1 |

| | | | | | |
|---|---|---|---|---|---|
| **26.** | *P. dicoccos* | rbcL | 99.66 | *Psydrax obovata* | KY378666.1 |
| | | Atp | 99.80 | *P. obovata* | KY378666.1 |
| | | PsbK | 95.73 | *P. obovata* | KY378666.1 |
| | | RpoB | 99.33 | *P. obovata* | KY378666.1 |
| | | PsbA-trnH1 | 98.04 | *P. obovata* | KY378666.1 |
| | | PsbA-trnH2 | 98.04 | *Psydrax dicoccos* | HQ415552.1 |
| | | Overall | 98.66 | *Psydrax obovata* | KY378666.1 |
| **27.** | *T. agumbensis* | RbcL1 | 96.59 | *Psychotria serpens* | LC692567.1 |
| | | Atp | 95.72 | *P. serpens* | NC_069807.1 |
| | | PsbA-trnH2 | 93.01 | *Psychotria* sp. | KF676320.1 |
| | | PsbK | 94.12 | *Psychotria kirkii* | KY378696.1 |
| | | RPB2 | 93.90 | *Bouvardia glaberrima* | DQ358891.1 |
| | | RpoB | 98.76 | *Psychotria rubra* | MZ958829.1 |
| | | Cn_psbZ-trnfM | 95.59 | *Psychotria rubra* | MZ958829.1 |
| | | Overall | 99.85 | *Psychotria rubra* | MZ958829.1 |
| **Passifloraceae** | | | | | |
| **28.** | *A. hondala* | RbcL1 | 98.97 | *Adenia* sp. | KF724302.1 |
| | | PsbAtrnH1 | 98.56 | *Adenia hondala* | MN167476.1 |
| | | Atp | 95.99 | *Adenia mannii* | NC_043791.1 |
| | | psbZ_trnfM | 92.59 | *A. mannii* | NC_043791.1 |
| | | RpoB | 99.05 | *A. mannii* | NC_043791.1 |
| | | RpoB_trnCGAR | 95.52 | *A. mannii* | NC_043791.1 |
| | | trnHpsbA2 | 99 | *Adenia hondala* | N167476.1 |
| | | Overall | 98.97 | *Adenia mannii* | KF724302.1 |

Supplementary File 2. GenBank Accession ids of submitted sequences of selected forest species

| **Selected species** | **DNA barcoding markers** | **Accession No** |
|---|---|---|
| **Family** | | |
| **Apocynaceae** | | |
| *R. serpentiana* | RbcL1 | OR260919 |
| | psbA-trnH1 | OR260920 |
| | psbZ | OR260921 |
| | rpob | OR260922 |
| | trnH-psbA2 | OR260923 |
| | Atp | OR260924 |
| *T. heyneana* | trnH-psbA2 | OR260967 |
| | Atp | OR260968 |
| | Psbk | OR260969 |
| | rpoB | OR260970 |
| **Anacardiaceae** | | |
| *H. arnottiana* | RbcL1 | OR236160 |
| | psbA-trnH1 | OR236161 |
| | Atp | OR236162 |
| | psbK | OR236163 |

| | | |
|---|---|---|
| | psbZ_trnfM | OR236164 |
| | RPB2 | OR236165 |
| | rpoB | OR236166 |
| | RpoB_trnCGAR | OR236167 |
| | trnH-psbA2 | OR236168 |
| | ITS2 | OR233169 |
| *S. auriculata* | matK1 | OR260938 |
| | rpoC | OR260939 |
| | RbcL1 | OR260940 |
| | psbA-trnH1 | OR260941 |
| | psbK | OR260942 |
| | Rpob | OR260943 |
| | RpoB-trnCGAR | OR260944 |
| | ITS | |
| **Arecaceae** | | |
| *C. nagbettai* | RbcL1 | OR236180 |
| | matK1 | OR236181 |

| | | |
|---|---|---|
| | psbA-trnH1 | OR236182 |
| | Atp | OR236183 |
| | PsbK | OR236184 |
| | Cn_psbZ_trnfM | OR236185 |
| | RPB2 | OR236186 |
| | RpoB | OR236187 |
| | trnH-psbA2 | OR236188 |
| *A. wightii* | matK1 | OR236149 |
| | RbcL1 | OR236150 |
| | psbA-trnH1 | OR236151 |
| | Atp | OR236152 |
| | PRK | OR236153 |
| | PsbK | OR236154 |
| | PsbZ-trnfM | OR236155 |
| | RPB2 | OR236156 |
| | RpoB | OR236157 |

| | | |
|---|---|---|
| | RpoC | OR236158 |
| | RpoBtrnCGAR | OR236159 |
| **Bignoniaceae** | | |
| *O. indicum* | matK1 | OR245475 |
| | RbcL1 | OR245476 |
| | psbA-trnH1 | OR245477 |
| | PsbK | OR245478 |
| | PsbZ-trnfM | OR245479 |
| | RpoB | OR245480 |
| | RpoC | OR245481 |
| | trnH-psbA2 | OR245482 |
| | RbcL2 | OR245483 |
| | Matk2 | OR245484 |
| | ITS2 | OR260478 |
| **Balsaminaceae** | | |
| *I. mysorensis* | RbcL1 | OR245434 |
| | psbA-trnH1 | OR245435 |

| | | |
|---|---|---|
| | Atp | OR245436 |
| | PsbK | OR245437 |
| | RpoB | OR245438 |
| | trnH-psbA2 | OR245439 |
| *I. pulcherrima* | matK 1 | OR245424 |
| | RbcL1 | OR245425 |
| | psbA-trnH1 | OR245426 |
| | Atp | OR245427 |
| | PsbK | OR245428 |
| | PsbZ-trnfM | OR245429 |
| | RpoC | OR245430 |
| | RpoBtrnCGAR | OR245431 |
| | trnH-psbA2 | OR245432 |
| | RbcL2 | OR245433 |
| **Calophyllaceae** | | |
| *C. apetalum* | RbcL1 | OR236169 |
| | psbA-trnH1 | OR236170 |

| | | |
|---|---|---|
| | Atp | OR236171 |
| | PsbK | OR236172 |
| | RpoB | OR236173 |
| | psbZ-trnfM | OR236174 |
| | trnH-psbA2 | OR236175 |
| Clusiaceae | | |
| *G. indica* | ITS2 | OR234847 |
| | psbZ-trnfM | OR245411 |
| *G. wightii* | matK1 | OR245399 |
| | RbcL1 | OR24539400 |
| | psbA-trnH1 | OR24539401 |
| | Atp | OR24539402 |
| | PsbK | OR24539403 |
| | PsbZ-trnfM | OR24539404 |
| | RPB2 | OR24539405 |
| | RpoB | OR24539406 |
| | RpoC | OR24539407 |

| | | |
|---|---|---|
| | RpoBtrnCGAR | OR24539408 |
| | trnH-psbA2 | OR24539409 |
| | RbcL2 | OR24539410 |
| **Cycadaceae** | | |
| *C. circinalis* | psbA-trnH1 | OR236176 |
| | Atp | OR236177 |
| | PRK | OR236178 |
| | RpoB_trnCGAR | OR236179 |
| **Dipterocarpaceae** | | |
| *S. roxburghii* | RbcL1 | OR260945 |
| | psbA-trnH1 | OR260946 |
| | trnH-psbA2 | OR260947 |
| | PsbK | OR260948 |
| | RPB2 | OR260949 |
| | RpoB | OR260950 |
| *H. ponga* | RPB2 | OR260916 |
| | trnH-psbA2 | OR260917 |

| | | |
|---|---|---|
| | PsbK | OR260918 |
| | ITS2 | OR260466 |
| *D. indicus* | ITS2 | OR233147 |
| | matK1 | OR242234 |
| | RbcL1 | OR242235 |
| | PsbK | OR242236 |
| | RpoB | OR242237 |
| | RpoC | OR242238 |
| | rbcL2 | OR242239 |
| *H. parviflora* | psbA-trnH1 | OR245419 |
| | PsbK | OR245420 |
| | trnH-psbA2 | OR245421 |
| *V. chinensis* | ITS2 | OR260436 |
| *D. bourdiollon* | matK1 | OR242240 |
| | RbcL1 | OR242241 |

| | | |
|---|---|---|
| | psbA-trnH1 | OR242242 |
| | PsbK | OR242243 |
| | RpoB | OR242244 |
| | RpoC | OR242245 |
| | RpoB-trnCGAR | OR242246 |
| | trnH-psbA2 | OR242247 |
| | RbcL2 | OR242248 |
| *V. macrocarpa* | RbcL1 | OR260979 |
| | psbA-trnH1 | OR260980 |
| | PsbK | OR260981 |
| | trnH-psbA2 | OR260982 |
| **Ebenaceae** | | |
| *D. paniculata* | RpoC | OR242232 |
| | RbcL1 | OR242233 |
| *D. crassiflora* | Md_ITS | OR233146 |
| | RbcL1 | OR238343 |
| | matK1 | OR238344 |

| | | |
|---|---|---|
| | psbA-trnH1 | OR238345 |
| | Atp | OR238346 |
| | PsbK | OR238347 |
| | PsbZ-trnfM | OR238348 |
| | RPB2 | OR238349 |
| | RpoB | OR238350 |
| | RpoB_trnCGAR | OR238351 |
| | trnH-psbA2 | OR238334 |
| **Fabaceae** | | |
| *D. latifolia* | ITS2 | OR233145 |
| | RbcL1 | OR238335 |
| | psbA-trnH1 | OR238336 |
| | Atp | OR238337 |
| | PsbZ-trnfM | OR238338 |
| | RPB2 | OR238339 |
| | RpoB | OR238340 |
| | RpoB_trnCGAR | OR238341 |

| | | |
|---|---|---|
| | trnH-psbA2 | OR238342 |
| *P. marsupium* | RbcL1 | OR245485 |
| | psbA-trnH1 | OR245486 |
| | RpoB | OR245487 |
| | trnH-psbA2 | OR245488 |
| *S. asoca* | RbcL1 | OR260933 |
| | psbA-trnH1 | OR260934 |
| | RPB2 | OR260935 |
| | RpoB | OR260936 |
| | trnH-psbA2 | OR260937 |
| *K. pinnatum* | trnH-psbA2 | OR245440 |
| | RpoB | OR245441 |
| | RpoB_trnCGAR | OR245442 |
| *P. santalinus* | RbcL1 | OR245492 |
| | psbA-trnH1 | OR245493 |
| | Atp | OR245494 |
| | RpoB | OR245495 |

| | | |
|---|---|---|
| | trnH-psbA2 | OR245496 |
| **Flacourtiaceae** | | |
| *H. macrocarpa* | RbcL1 | OR245422 |
| | RpoC | OR245423 |
| **Lauraceae** | | |
| *C. riparium* | matK1 | OR238322 |
| | RbcL1 | OR238323 |
| | Atp | OR238324 |
| | psbK | OR238325 |
| | psbZ-trnfM | OR238326 |
| | RPB2 | OR238327 |
| | rpoB | OR238328 |
| | rpoC | OR238329 |
| | psbA-trnH1 | OR238330 |
| | matK2 | OR238331 |
| | trnH-psbA2 | OR238332 |
| | RbcL2 | OR238333 |

| | | |
|---|---|---|
| **Primulaceae** | | |
| *E. ribes* | ITS2 | OR234841 |
| | matK1 | OR245388 |
| | RbcL1 | OR245389 |
| | psbA-trnH1 | OR245390 |
| | Atp | OR245391 |
| | Psbk | OR245392 |
| | psbZ-trnfM | OR245393 |
| | rpoB | OR245394 |
| | rpoC | OR245395 |
| | RpoB-trnCGAR | OR245395 |
| | trnH-psbA2 | OR245397 |
| | RbcL2 | OR245398 |
| **Malvaceae** | | |
| *P. reticulatum* | RbcL1 | OR245489 |
| | psbK | OR245490 |
| | rpoB | OR245491 |

| | | |
|---|---|---|
| *S. occidentale* | matK1 | OR260951 |
| | RbcL1 | OR260952 |
| | psbA-trnH1 | OR260953 |
| | Atp | OR260954 |
| | psbK | OR260955 |
| | psbZ-trnfM | OR260956 |
| | RPB2 | OR260957 |
| | rpoB | OR260958 |
| | rpoC | OR260959 |
| | RpoB-trnCGAR | OR260960 |
| | trnH-psbA2 | OR260961 |
| | RbcL2 | OR260962 |
| *S. travancoricum* | RbcL1 | OR260963 |
| | psbZ-trnfM | OR260964 |
| | rpoB | OR260965 |
| | trnH-psbA2 | OR260966 |
| **Meliaceae** | | |

| | | |
|---|---|---|
| *D. malabaricum* | matK1 | OR242249 |
| | Atp | OR242250 |
| | rpoC | OR242251 |
| | RbcL1 | OR242252 |
| | psbK | OR242253 |
| | rpoB | OR242254 |
| | RbcL2 | OR242255 |
| **Myristicaceae** | | |
| *G. canarica* | matK1 | OR245412 |
| | psbA-trnH1 | OR245413 |
| | psbK | OR245414 |
| | psbZ-trnfM | OR245415 |
| | Rpob | OR245416 |
| | rpoc | OR245417 |
| | trnH-psbA2 | OR245418 |
| *M. malabarica* | RbcL1 | OR245465 - |
| | matK1 | OR245466 |

| | | |
|---|---|---|
| | psbA-trnH1 | OR245467 |
| | Atp | OR245468 |
| | psbK | OR245469 |
| | psbZ-trnfM | OR245470 |
| | RPB2 | OR245471 |
| | RpoB | OR245472 |
| | RpoB-trnCGAR | OR245473 |
| | trnH-psbA2 | OR245474 |
| *M. beddomei* | RbcL1 | OR245458 |
| | psbA-trnH1 | OR245459 |
| | Atp | OR245460 |
| | psbK | OR245461 |
| | psbZ-trnfM | OR245462 |
| | Rpoc | OR245463 |
| | trnH-psbA2 | OR245464 |
| *K. attenuata* | RbcL1 | OR236143 |

| | | |
|---|---|---|
| | psbA-trnH1 | OR236144 |
| | Atp | OR236145 |
| | psbK | OR236146 |
| | psbZ | OR236147 |
| | trnH-psbA2 | OR236148 |
| **Sapotaceae** | | |
| *M. bourdillonii* | RbcL1 | OR245443 - |
| | psbA-trnH1 | OR245444 |
| | psbK | OR245445 |
| | Atp | OR245446 |
| | rpoC | OR245447 |
| | trnH-psbA2 | OR245448 |
| | RbcL2 | OR245449 |
| *M. insignis* | RbcL1 | OR245450 |
| | psbA-trnH1 | OR245451 |
| | Atp | OR245452 |
| | psbZ-trnfM | OR245453 |

| | | |
|---|---|---|
| | RPB2 | OR245454 |
| | RpoB | OR245455 |
| | RpoB-trnCGAR | OR245456 |
| | trnH-psbA2 | OR245457 |
| | **Rubiaceae** | |
| *O. missionis* | RbcL1 | OR188689 |
| | psbA-trnH1 | OR188690 |
| | Atp | OR228661 |
| | PsbK | OR228663 |
| | RPB2 | OR228650 |
| | RpoB | OR228657 |
| | trnH-psbA2 | OR228668 |
| | Ga_RpoB-trnCGAR | OR228656 |
| *P. dicoccos* | RbcL1 | OR260911 |
| | trnH-psbA2 | OR260912 |
| | Atp | OR260913 |

| | | |
|---|---|---|
| | psbK | OR260914 |
| | Rpob | OR260915 |
| *T. agumbensis* | RbcL1 | OR260971 |
| | psbA-trnH1 | OR260972 |
| | Atp | OR260973 |
| | psbK | OR260974 |
| | psbZ-trnfM | OR260975 |
| | RPB2 | OR260976 |
| | rpoB | OR260977 |
| | trnH-psbA2 | OR260977 |
| **Santalaceae** | | |
| *S. album* | RbcL1 | OR260925 |
| | psbA-trnH1 | OR260926 |
| | Atp | OR260927 |
| | psbK | OR260928 |
| | psbZ-trnfM | OR260929 |
| | rpoB | OR260930 |

| | | |
|---|---|---|
| | trnCGA | OR260931 |
| | trnH-psbA2 | OR260932 |
| **Passifloraceae** | | |
| *A. hondala* | RbcL1 | OR228651 |
| | psbA-trnH1 | OR228653 |
| | Atp | OR228660 |
| | psbK | OR228662 |
| | psbZ_trnfM | OR228664 |
| | rpoB | OR228658 |
| | trnH-psbA2 | OR228667 |
| | RpoB_trnCGAR | OR228655 |
